# MDM2 acts as a timer reporting the length of mitosis

**DOI:** 10.1101/2023.05.26.542398

**Authors:** Luke J. Fulcher, Tomoaki Sobajima, Caleb Batley, Ian Gibbs-Seymour, Francis A. Barr

## Abstract

Delays in mitosis trigger p53-dependent arrest in G1 of the following cell cycle, enabling cells to respond to changes that would otherwise promote chromosome instability and aneuploidy ^1–10^. We find that MDM2, the p53 ubiquitin ligase, is a key component of the timer mechanism triggering G1 arrest in response to prolonged mitosis. This timer function arises because MDM2 has a short half-life and ongoing protein synthesis is therefore necessary to maintain its steady-state concentration. Due to the attenuation of protein synthesis in mitosis, the amount of MDM2 gradually falls during mitosis, but normally remains above a critical threshold for p53 regulation at the onset of G1. When mitosis is extended by prolonged spindle assembly checkpoint activation, the amount of MDM2 drops below this threshold, stabilising p53. Subsequent p53-dependent p21 accumulation in the following G1 then channels cells into a prolonged cell cycle arrest, whereas abrogation of the response in p53-deficient cells allows them to bypass this crucial defence mechanism.

## Main

Chromosome instability, aneuploidy, the removal of centrosomes, or anti-mitotic drugs targeting the microtubule cytoskeleton delay progression through mitosis, triggering a p53 and p21 dependent cell cycle arrest in G1 thought to prevent proliferation of damaged cells ^1–6^. This response is lost in cancer cells in which p53 has become inactivated by mutation or other mechanisms, including expression of viral oncoproteins ^7–10^. Crucially, G1 cell cycle arrest following prolonged mitosis occurs even in the absence of detectable DNA damage, suggesting it has a different cause, proposed to be a direct consequence of the increased time spent in mitosis ^1–10^. Therefore, we asked if variation in the length of mitosis, inherent in the stochastic search-capture process underpinning chromosome alignment even in the absence of any perturbation, influences the behaviour of untransformed diploid cells with wild-type p53 (p53^WT^) in the ensuing G1. To do this, we used fluorescence imaging and single cell tracking of telomerase-immortalised retinal pigmented epithelium cells (hTERT-RPE1) stably expressing a FUCCI reporter ^11^. For individual cells under normal growth conditions, we measured the length of mitosis and subsequent G1 duration, and recorded whether the cell entered S-phase (Fig. 1a-1c). The majority of p53^WT^ cells completed mitosis in under 60 minutes, with a mean time of 50.1 ± 10.8 minutes (Fig. 1b), and then passed through G1 and entered S-phase in 10.2 ± 0.3 hours for cells spending 40-49 minutes in mitosis or 12.4 ± 0.6 hours, slightly longer, for cells spending 50-59 minutes in mitosis (Fig. 1a and 1c). By contrast, p53^WT^ cells which spent more than 60 minutes in mitosis arrested in G1 for at least 29.1 ± 1.7 hours without any notable increase in cell death (Fig. 1a and 1c). For hTERT-RPE1 p53 knockout (p53^KO^) cells, mean time in mitosis was 51.5 ± 12.5 minutes, not significantly different to p53^WT^ cells (Fig. 1b), showing p53 does not play a direct role in mitotic timing. However, unlike p53^WT^ cells, increased G1 length and cell cycle arrest were not observed in p53^KO^ cells spending longer than 60 minutes in mitosis (Fig. 1a and 1c). Thus, there is a threshold time in mitosis beyond which cells respond in a p53-dependent process and undergo cell cycle arrest in the subsequent G1.

**Fig. 1.**
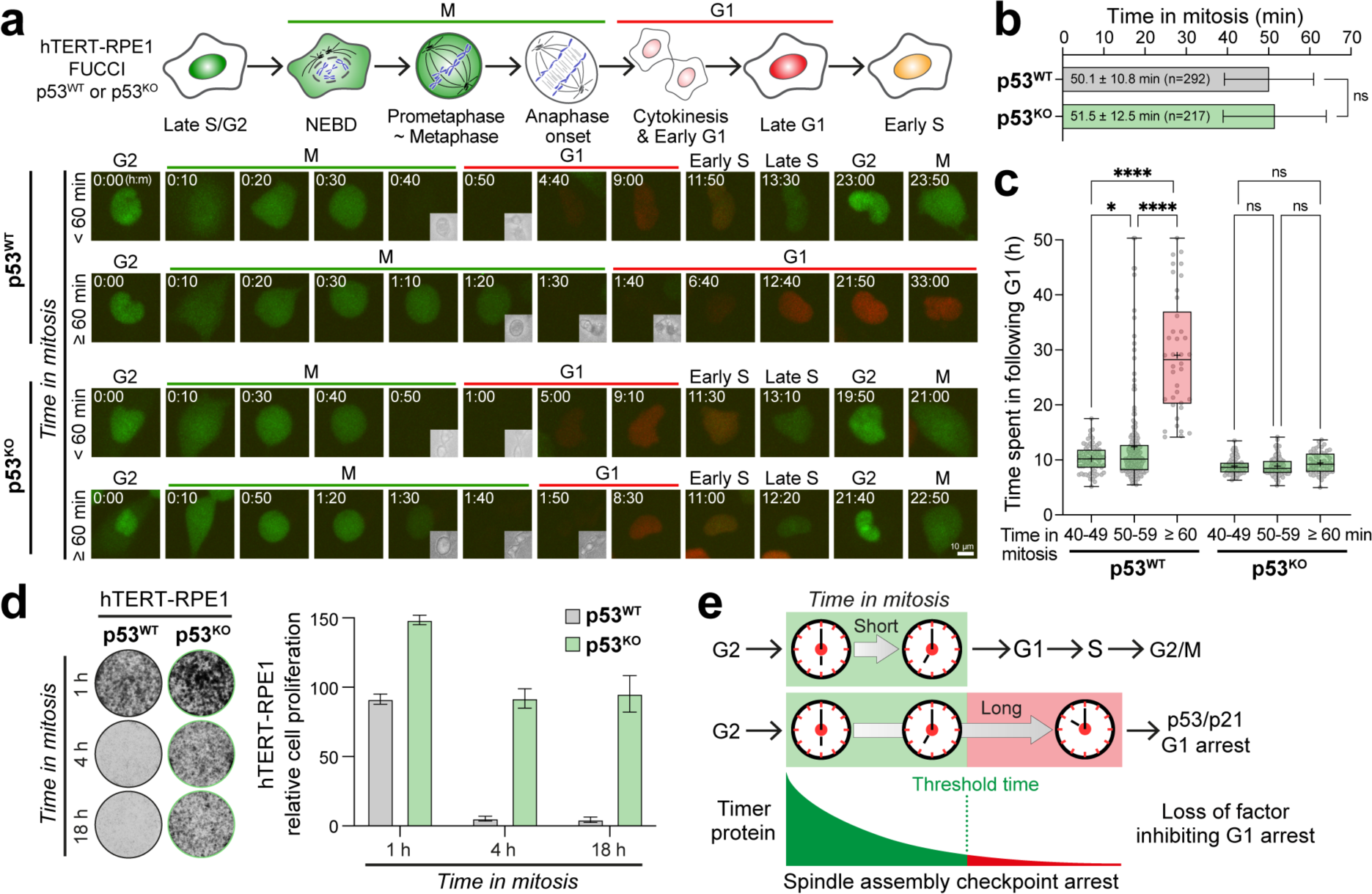
Stochastic variation in mitosis beyond a defined time threshold triggers a p53-dependent cell cycle arrest in the ensuing G1. **a.** A schematic of the hTERT-RPE1 FUCCI cell line with representative images of asynchronous cultures of hTERT-RPE1 p53^WT^ and p53^KO^ FUCCI cells entering and exiting mitosis, and progressing into the following cell cycle, grouped according to time in mitosis. G2 cells were identified, then tracked and imaged every 10 minutes up to 60 h post-mitosis. Insets show anaphase onset and cytokinesis. **b.** The mean length of mitosis is plotted as mean ± SD. **c.** The duration of G1 for p53^WT^ cells spending 40-49 (n=69), 50-59 (n=185) or ≥ 60 minutes (n=38) in mitosis, or for p53^KO^ cells spending 40-49 (n=65), 50-59 (n=100) or ≥ 60 minutes (n=52) is shown in a box and whiskers plot with the mean G1 length (+). **d.** hTERT-RPE1 p53^WT^ or p53^KO^ cells were arrested in mitosis for 1, 4 or 18 h with 25 ng/ml nocodazole. Mitotic cells were harvested, washed-out from nocodazole, and 5000 cells plated per well. After 5 days cell proliferation was measured by crystal violet staining (mean ± SEM, n=5). Representative stained wells are shown. **e.** A mitotic timer pathway triggers p53-dependent arrest in G1 following delayed mitosis. Loss of a factor inhibiting G1 arrest could act as a sensor for the length of mitosis in this timer pathway.

To more directly test for a link between time in mitosis and G1 cell cycle arrest, we used transient exposure to a low dose of the microtubule poison nocodazole to perturb chromosome alignment and activate the spindle assembly checkpoint for up to 1, 4 or 18 h, similar to the conditions used in earlier studies ^1–10^. Checkpoint-arrested mitotic cells were collected and then replated in fresh growth medium lacking nocodazole and tested for proliferation. It is important to note that the low dose of nocodazole arrests cells in mitosis with complete spindles and few unaligned chromosomes (Fig. S1a). When the drug is washed out, these chromosomes efficiently align and cells rapidly progress into anaphase, without chromosome segregation errors, and then into G1 to give normal shaped nuclei (Fig. S1b). Cell proliferation was still observed after 1 h in mitosis (Fig. 1d), consistent with the FUCCI imaging assays showing this time remains below the threshold needed to trigger G1 arrest. Prolonging mitosis from 1 h to 4 h or 18 h by activating the spindle assembly checkpoint resulted in cell cycle arrest and a cessation of cell proliferation in hTERT-RPE1 p53^WT^ cells (Fig. 1d). In contrast, an hTERT-RPE1 p53^KO^ cell line showed normal proliferation after 1, 4 or 18 h mitotic delay (Fig. 1d), demonstrating the cell cycle arrest is mediated by a p53-dependent pathway. This p53-dependent reduction in proliferation of asynchronously growing cells with extended mitosis (Fig. 1a-1c), or cells with spindle checkpoint delayed mitosis (Fig. 1d), was not associated with DNA damage in the arrested G1 cells (Fig. S2a-S2c), suggesting it has another cause. We therefore explored the idea that the detection of delays in mitosis is linked to a biochemical reaction consuming a key regulatory component of the p53 pathway, which thus acts as a timer reporting the length of mitosis (Fig. 1e). This led us to consider the role of MDM2, the p53 E3 ubiquitin-ligase ^12–14^.

### The short half-life and attenuation of MDM2 synthesis in mitosis create a potential timer

The levels of p53 are regulated by ongoing synthesis and proteasomal destruction triggered by the MDM2 E3 ubiquitin-ligase. DNA damage and other stress signals mediated by conserved signalling pathways inhibit MDM2 activity towards p53 ^15^. This results in p53 stabilization, an increase in the steady-state concentration of p53, and p53-dependent transcription of target genes, including MDM2, DNA repair proteins, cell cycle regulators such as the CDK-inhibitor p21, and, if DNA damage is not repaired, pro-apoptotic factors ^15^. Like p53, MDM2 stability is tightly regulated. MDM2 has a short half-life and is subject to autoubiquitination or ubiquitination by other E3 ubiquitin-ligases ^16,17^. We inferred that entry into mitosis would have important functional consequences for both MDM2 itself and MDM2 regulation of p53 at the onset of the following G1 (Fig. 2a). Upon entry into mitosis there is a general attenuation of both transcription and translation, until cells exit mitosis and re-enter G1, when transcription and protein synthesis resume ^18–20^. We therefore hypothesized that MDM2 synthesis would stop or become greatly reduced following mitotic entry, whereas its turnover would continue due to ongoing self-catalysed ubiquitination or ubiquitination by other E3 ubiquitin-ligases. On this basis, we predicted that early G1 cells arising from a normal length mitosis must retain sufficient MDM2 to regulate p53 normally and prevent p21-induced cell cycle arrest. However, if mitosis is prolonged and MDM2 concentration falls below this threshold, it would become limiting for p53 regulation early in the following G1, resulting in p53 stabilization and p21-dependent cell cycle arrest. Our hypothesis is that these are general properties of the MDM2-p53 pathway, providing a mechanism to monitor the length of mitosis, independent from the cause of the delay. While elements of this hypothesis have been discussed previously ^21,22^, to date there is no experimental evidence in support of this proposal. We therefore set out to test these ideas and whether MDM2 has the hallmarks predicted for the mitotic timer protein.

**Fig. 2.**
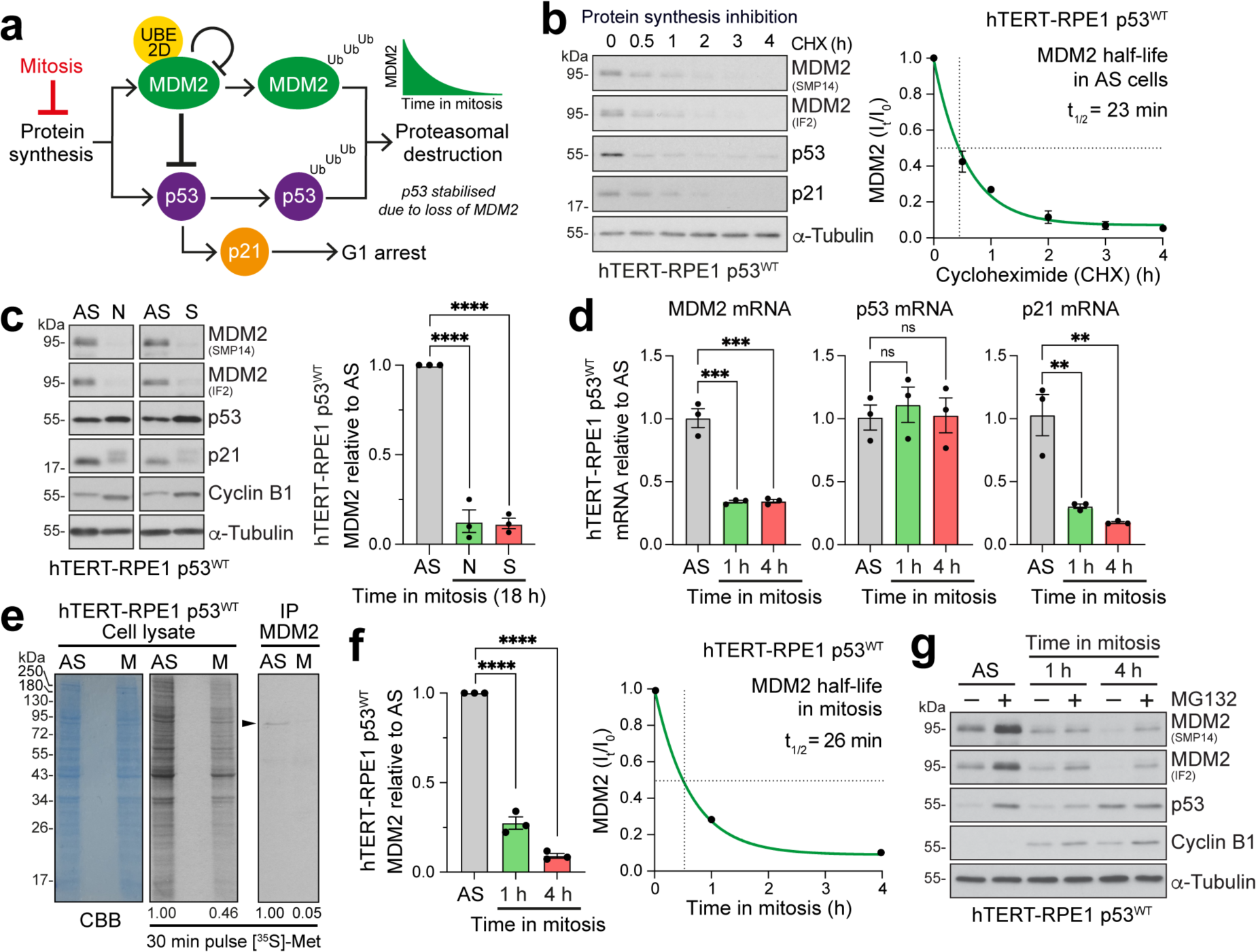
MDM2 synthesis but not turnover is attenuated in mitosis. **a.** A schematic depicting MDM2 regulation of p53, and the role of ongoing protein synthesis and proteolysis in this pathway. **b.** MDM2 half-life was measured in asynchronous hTERT-RPE1 p53^WT^ cells using Western blotting by adding 50 µg/ml cycloheximide (CHX) to block protein synthesis, and then collecting samples up to 4 h. MDM2 half-life is indicated in the graph (mean ± SEM, n=3), with example Western blots. **c.** The amount of MDM2 was measured by Western blotting in asynchronous (AS) or mitotic hTERT-RPE1 p53^WT^ cells, arrested in mitosis for 18 h with 100 ng/ml nocodazole (N) or 5 µM S-trityl L-cysteine (STLC, S) (mean ± SEM, n=3). **d.** RT-qPCR for the level of MDM2, p53 and p21 mRNAs in asynchronous cell cultures, or cells arrested in mitosis for 1 or 4 h with 25 ng/ml nocodazole (mean ± SEM, n=3). **e.** Cell lysates and MDM2 immunoprecipitations (IP) from asynchronous (AS) or mitotically arrested (M) hTERT-RPE1 p53^WT^ cells labelled for 30 minutes with [^35^S]-methionine. Numbers indicate the [^35^S]-methionine signal relative to the AS control. **f.** The amount of MDM2 was measured by Western blotting in asynchronous (AS) or mitotic hTERT-RPE1 p53^WT^ cells, arrested in mitosis for 1 or 4 h with 25 ng/ml nocodazole (mean ± SEM, n=3). Mitotic MDM2 half-life was calculated from the curve fit plotted on the right. **g.** As in f., except cells were treated ± 20 µM MG-132 to inhibit the proteasome.

First, we measured the stability of MDM2 in untransformed hTERT-RPE1 p53^WT^ cells, and a range of transformed human cancer cell lines, in both interphase and mitotic states. MDM2 and p53 expression was confirmed for all cell lines in asynchronous culture with two different MDM2 antibodies (Fig. S3a). HeLa cells have much lower levels of MDM2 and p53 than other cell lines due to expression of the HPV E6 protein which targets p53 for destruction, and because MDM2 is a transcriptional target of p53 ^23,24^. Using cycloheximide to block new protein synthesis, we found that endogenous MDM2 has a half-life of ∼30 minutes in all of the cell lines tested, except HeLa where it is shorter due to the presence of HPV E6 (Fig. 2b and S4). We then arrested these different cell lines in mitosis with two different anti-mitotic agents (nocodazole or the Eg5 kinesin inhibitor STLC) and measured the amount of MDM2 by Western blotting. These approaches revealed that MDM2 falls to very low levels during an 18 h mitotic arrest in all cell lines, regardless of which agent was used to trigger the arrest (Fig. 2c, S3b and S3c). In contrast, p53 was stable in a prolonged mitotic arrest in all cell lines except HeLa, consistent with the idea that its destruction requires MDM2 (Fig. 2c and S3b). Compared to the other cell lines, HeLa cells had very low levels of p53 due to HPV E6 expression, which fell below the detection limit in the mitotic arrest samples (Fig. S3b). To explain the decrease in MDM2 we next explored the roles of new protein synthesis and turnover. Using RT-qPCR, we measured the levels of MDM2 mRNA relative to a housekeeping gene GAPDH in asynchronous cell cultures and cells arrested in mitosis for different lengths of time (Fig. 2d, S5a-S5c). This revealed that, like the level of MDM2 protein, MDM2 mRNA drops rapidly following entry into mitosis within 1 h and does not recover up to 18 h of mitotic arrest (Fig. 2d, Fig. S5c and S5d). Similar behaviour was observed for p21 mRNA, although the protein, albeit modified in mitosis, was stable (Fig. 2c and 2d, Fig. S5c and S5d). This means that new p21 synthesis in G1 following a mitotic delay would require p53-dependent transcription of the mRNA ^25,26^. By contrast, p53 mRNA and protein were largely unaltered in abundance in mitotically arrested cells compared to asynchronous cells (Fig. 2c and 2d, Fig. S5c and S5d). We next tested whether MDM2 protein synthesis is altered in mitosis using 30-minute pulse-labelling of asynchronous and mitotic cells with [^35^S]-methionine. The global level of protein synthesis was reduced over two-fold in mitotic cells compared to an asynchronous culture, whereas MDM2 synthesis decreased by over 95% (Fig. 2e). We then investigated the half-life of MDM2 in mitotic hTERT-RPE1 cells. MDM2 levels declined dependent on the length of time in mitosis, showing a robust decrease from 1 to 4 h of mitotic delay with an estimated half-life for MDM2 of ∼26 min (Fig. 2f). Addition of the proteasome inhibitor MG132 prevented MDM2 destruction after 4 h of mitotic delay (Fig. 2g). Thus, reduction in the level of MDM2 mRNA and global attenuation of protein synthesis, combined with the short half-life of the MDM2 protein, can explain how the amount of MDM2 decreases in mitotic cells through a ubiquitination-dependent pathway.

### MDM2 catalyses its own destruction in a UBE2D-dependent manner

The mechanism of MDM2 turnover in mitosis is a critical component of the proposed timer function. As already shown, this is likely to be via a ubiquitination-dependent pathway (Fig. 2g). MDM2 has been shown to undergo both self-catalysed ubiquitination, as well as ubiquitination by other E3 ubiquitin ligases ^16,17^. We favoured a mechanism whereby MDM2 catalyses its own destruction (Fig. 3a), since this renders the rate of MDM2 turnover less dependent on other factors including MDM2 concentration, assuming the availability of charged E2-ubiquitin conjugates is not limiting. To investigate this potential self-catalysed ubiquitination mechanism, we asked which domains of MDM2 are required for its turnover. Using cycloheximide to inhibit protein synthesis, we have mapped MDM2 turnover to the C-terminal RING domain responsible for E3-ubiquitin ligase activity towards p53 (Fig. 3b). Published structure-function data enabled us to generate specific I440A and R479A point mutants on the surface of the RING domain that prevent E2-ubiquitin binding, but do not alter the structure ^27^. These mutants show stabilization compared to wild-type MDM2 (Fig. 3b). In further support of the idea that E2-binding is necessary for MDM2 destruction, combined depletion of UBE2D2 and UBE2D3, the MDM2 E2 enzymes ^28^, resulted in a pronounced increase in the amount of MDM2 and stabilisation of MDM2 when protein synthesis was inhibited in asynchronous cells (Fig. 3c). Moreover, in agreement with the idea that MDM2 destruction is ubiquitin dependent, inhibition of the proteasome stabilised MDM2 and resulted in a rise of concentration over time (Fig. 3d). Combined depletion of UBE2D2 and UBE2D3 also resulted in pronounced increase in MDM2 stability during mitosis (Fig. 3e). UBE2D2 and UBE2D3 are also required for the activity of MDM2 towards p53, and thus their depletion also stabilises p53 (Fig. 3e) and triggers a G1 arrest (Fig. 3f and 3g). To act as a timer, it is crucial that MDM2 isn’t destroyed to completion in every cell cycle regardless of the length of mitosis. It was therefore important to establish if either of the two major cell cycle ubiquitin ligases associated with mitosis and G1, the anaphase promoting complex/cyclosome (APC/C) and Skp1-Cullin-F-box (SCF) play a role in MDM2 destruction ^29^. MDM2 turnover was not altered by depletion of either the APC/C co-activator cell division cycle 20 homologue (CDC20) or both isoforms of the β-transducin-repeat containing protein (β-TrCP) a major substrate binding subunit for SCF, or by addition of the APC/C inhibitor proTAME (Fig. S6a-S6d).

**Fig. 3.**
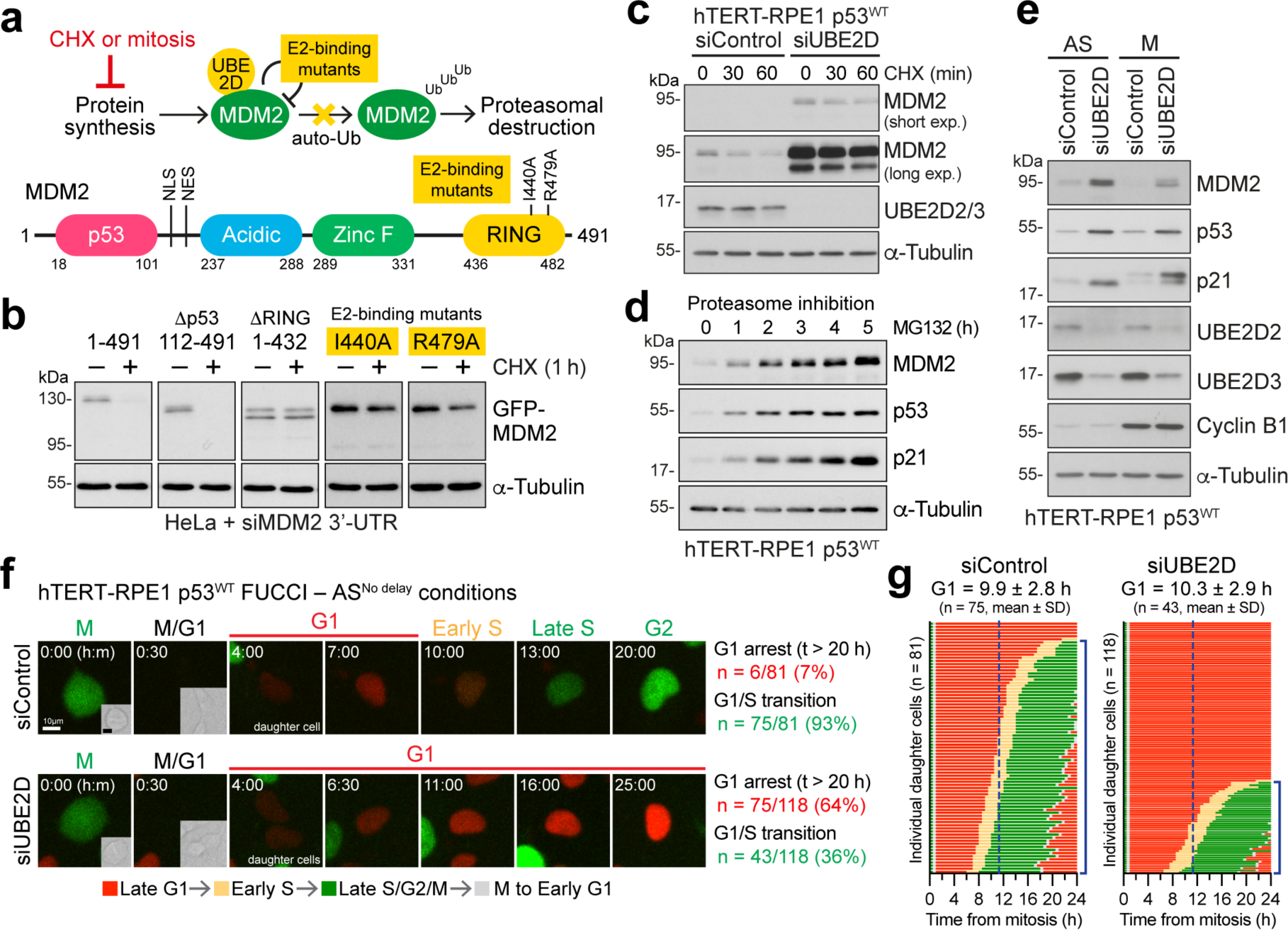
MDM2 catalyses its own destruction in a process requiring the RING domain and UBE2D. **a.** A schematic depicting regulation of MDM2 by auto-ubiquitination (top). The domain structure of MDM2 is cartooned with two E2-binding mutants (bottom). **b.** The stability of full-length GFP-MDM2 (1-491), and deletion or point mutant constructs were tested in HeLa cells depleted of endogenous MDM2 using a 3’-targeting siRNA, using 50 µg/ml cycloheximide (CHX) to block protein synthesis for 1 h (+). Solvent lacking cycloheximide was used as a control (–). **c.** The stability of endogenous MDM2 was tested in hTERT-RPE1 p53^WT^ cells, depleted of the MDM2 E2 enzymes UBE2D2/3 (siUBE2D), using 50 µg/ml CHX up to 1 h. A non-targeting siRNA was used as the control. **d.** The amount of MDM2 was measured by Western blotting in hTERT-RPE1 p53^WT^ cells treated for 0 to 5 h with 20 µM of the proteasome inhibitor MG132. **e.** Endogenous MDM2 stability was measured in hTERT-RPE1 p53^WT^ cells depleted of the MDM2 E2 enzymes UBE2D2/3 (siUBE2D) for 30 h in asynchronous culture (AS) or after arrest in mitosis (M). Non-targeting siRNA was used as a control. **f.** Representative images of control or UBE2D-depleted hTERT-RPE1 p53^WT^ FUCCI cells exiting mitosis and entering the following cell cycle. Mitotic cells from asynchronous culture (AS^NoDelay^) were tracked and imaged up to 60 h post-mitosis. The number and proportion of cells arresting in G1 and cells entering S-phase are shown. **g.** Single cell traces of hTERT-RPE1 p53^WT^ FUCCI cells treated as in panel f. passing from mitosis into the following cell cycle. The duration of G1 to cells entering S-phase is shown for all conditions (mean ± SD, sample sizes indicated in the figure). The dotted line in each panel of traces indicates the mean length of G1.

Together, these results provide support for the notion that MDM2 catalyses its own destruction in mitosis, and its half-life is an intrinsic property independent of the canonical cell cycle ubiquitin-ligases. However, because of the dual role of UBE2D as an E2 for both p53 and MDM2 itself, its manipulation does not allow us to fully test the role of MDM2 concentration as a key factor in the mitotic timer pathway.

### MDM2 destruction in mitosis results in p53 stabilisation, p21-induction and cell cycle arrest

Testing the proposed mitotic timer mechanism is complicated, since removal or manipulation of MDM2 activity using standard genetic methods will cause p53 stabilization and cell cycle arrest, independent from mitosis. We therefore needed to specifically target MDM2 within mitosis, and uncouple its stability from the length of time spent in mitosis. To selectively modulate MDM2 stability, we used the MDM2 PROteolysis-TArgeting Chimera (PROTAC) reagent MD-224 ^30^, which combines a derivative of the MDM2-binding drug Nutlin-3a with a ligand for the Cereblon-Cullin 4A-RBX1 E3 ubiquitin-ligase (Fig. S7a). In asynchronous cultures of hTERT-RPE1 p53^WT^ cells we observed that MD-224 caused efficient Cereblon and proteasome-dependent destruction of MDM2 after 1 h, and that its removal resulted in the rapid re-accumulation of MDM2 to its normal steady-state level within 1 h (Fig. S7b-S7d). The latter result was important since we needed to add MD-224 in mitosis, and then wash it out before cells exit mitosis and enter G1. Confirming the anticipated functional consequences of MDM2 destruction under these conditions, we observed p53 stabilisation, and after a short delay, an increase in p21 during the MD-224 washout period (Fig. S7c). In hTERT-RPE1 p53^KO^ cells, MD-224 resulted in rapid MDM2 destruction, but the amount of MDM2 did not recover rapidly once MD-224 was washed out (Fig. S7c), since MDM2, like p21, is a transcriptional target of p53 ^24^. Importantly, we confirmed that MD-224 was also active in cells arrested in mitosis, and promoted efficient MDM2 destruction within 1 h in both hTERT-RPE1 p53^WT^ and p53^KO^ cells (Fig. S7e). Compared to asynchronous cell cultures where most cells are in G1, S-phase or G2, the amount of MDM2 was reduced even with 1 h mitotic delay in the absence of MD-224 (Fig. S7e and S7f). This is consistent with the results in Figure 2 showing loss of MDM2 in prolonged mitosis. As expected, given the mode of action of MD-224, MDM2 destruction was prevented when mitotic cells were pre-incubated with the proteasome inhibitor MG132 (Fig. S7f).

These results enabled us to test the idea that the stability of MDM2 is a key component of the mitotic timer. For this purpose, we combined a short 1 h or long 4 h spindle checkpoint arrest with MD-224 to destabilise MDM2 in hTERT-RPE1 p53^WT^ and p53^KO^ cells (Fig. 4a). Biochemical analysis of these cells confirmed that MDM2 was still present after a 1 h mitotic delay, albeit reduced in amount in comparison to asynchronous cells (Fig. 4b, lanes 1 and 9; Fig. 4c), and was further reduced to undetectable levels when MD-224 was added during that 1 h delay (Fig. 4b, lanes 1 and 2; Fig. 4c). After a 4 h delay, MDM2 was strongly reduced even in the absence of MD-224, and was further reduced to undetectable levels if MD-224 was present (Fig. 4b, lanes 1, 5 and 6; Fig. 4c). In p53^WT^ cells, p53 and p21 increased significantly in the G1 following a 1 h mitotic delay, when MDM2 was destabilised in mitosis by MD-224, to the levels seen after a 4 h mitotic delay (Fig. 4b, compare lanes 3-4 with lanes 7-8; Fig. 4d and 4e). Crucially, a short 1 h mitotic delay in the presence but not the absence of MD-224 triggered a cell cycle arrest and loss of proliferation in p53^WT^ cells (Fig. 4f). A long 4 h mitotic delay triggered a cell cycle arrest and loss of proliferation in both the presence and absence of MD-224 (Fig. 4f), suggesting that after 4 hours in mitosis MDM2 concentration had already fallen below the threshold needed to prevent p53 stabilisation and induction of p21.

**Fig. 4.**
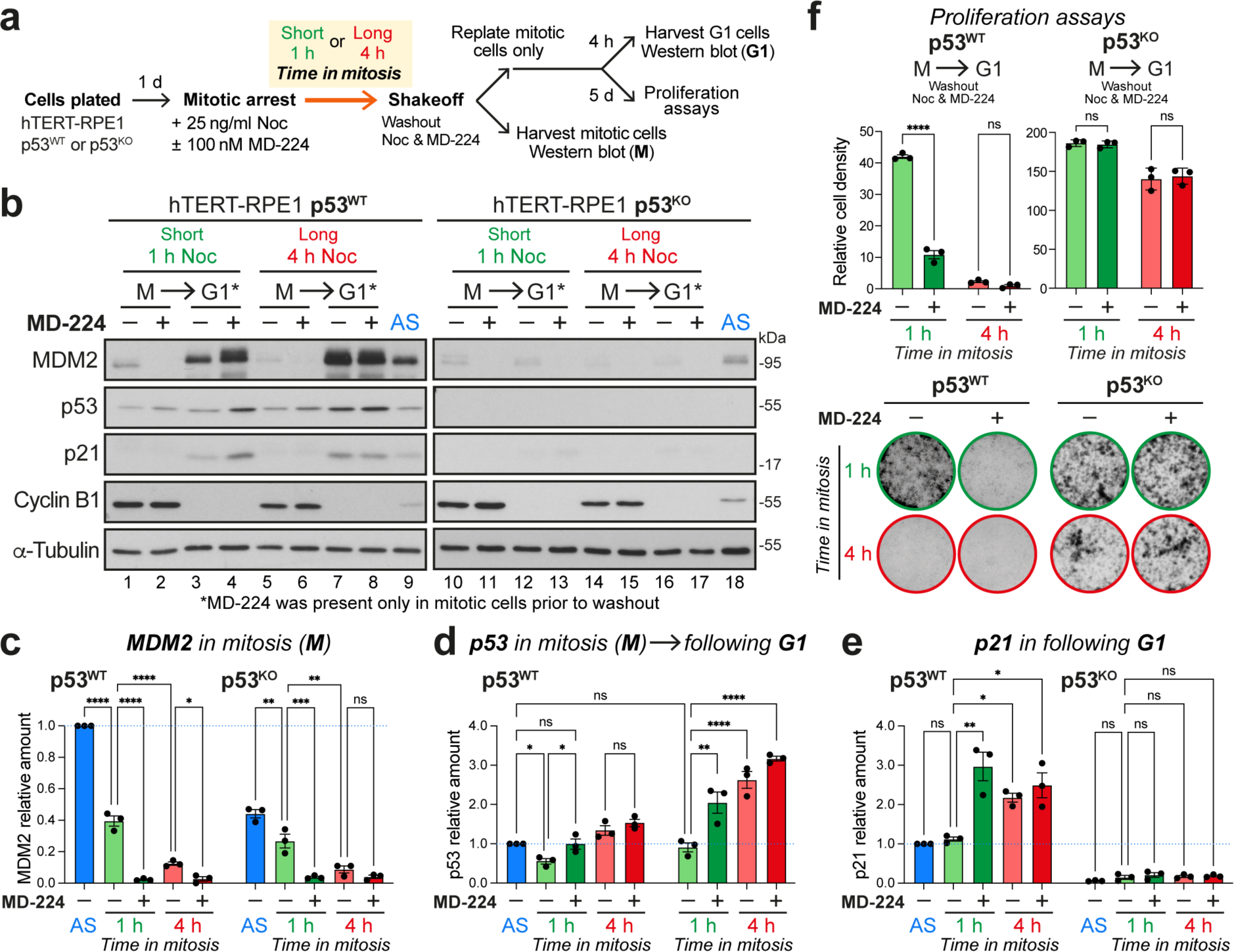
Mitotic delays or selective MDM2 destabilisation in mitosis result in p53-dependent p21-induction and reduced cell proliferation. **a.** A plan of the experimental design used for b-f. **b.** Western blot of cyclin B1 positive mitotic (M) and cyclin B1 negative (G1) hTERT-RPE1 p53^WT^ (lanes 1-8) or p53^KO^ cells (lanes 10-17) arrested for either 1 h or 4 h in mitosis with 25 ng/ml nocodazole, in the absence (–) or presence (+) of 100 nM MD-224 for the final hour. Samples from asynchronous hTERT-RPE1 p53^WT^ (AS, lane 9) and p53^KO^ cells (AS, lane 18) were loaded to indicate the steady-state level of MDM2, p53 and p21 prior to entry into mitosis. **c-e.** Graphs showing the relationship between mitotic delay and the amount of MDM2 (**c**) and p53 (**d**) in mitosis and p53 (**d**) and p21 (**e**) in the following G1 in p53^WT^ and p53^KO^ hTERT-RPE1 cells (mean ± SEM, n=3). Asynchronous hTERT-RPE1 p53^WT^ cells (AS) were used as a normalisation control for the graphs to allow direct comparison of the steady-state levels of MDM2, p53 and p21 in p53^WT^ and p53^KO^ cells. **f.** Cell proliferation was measured for both hTERT-RPE1 p53^WT^ or p53^KO^ mitotic cells treated as in b, using colony formation assays (mean ± SEM, n=3).

Matching this behaviour, p53 was stabilised in the subsequent G1, leading to upregulation of its transcriptional targets, including MDM2 (compare Fig. 4b, lanes 3-4 with 7-8, see also Fig. 4c). This behaviour confirmed that MD-224 had been efficiently removed prior to G1 entry, supporting the view that its effects are due to the destabilisation of MDM2 in mitosis. Confirming the dependence on p53 for the G1 arrest, induction of p21 and reduced cell proliferation were not observed with hTERT-RPE1 p53^KO^ cells for any of these conditions (Fig. 4b lanes 10-18, and Fig. 4e-4f). Note that because MDM2 is a transcriptional target of p53 it shows reduced steady-state levels in p53^KO^ cells compared to p53^WT^ cells (compare Fig. 4b, lane 9 AS p53^WT^ with lane 18 AS p53^KO^, see also Fig. 4c). The amount of MDM2 remaining at the end of mitosis is therefore an important determinant for whether cells undergo p53-dependent arrest in the following G1.

### Defining the MDM2 threshold in mitosis for p21 induction and robust cell cycle arrest in G1

Next, the cell cycle path taken by individual cells undergoing normal length or delayed mitosis in the presence and absence of MD-224 was analysed. Single cell tracing of hTERT-RPE1 p53^WT^ FUCCI cells was used to follow mitotic exit, the length of G1, and entry into S-phase (Fig. S8a and S8b) ^11^. For these experiments, mitotic cells were tracked after nocodazole washout following a short or long mitotic delay, or from asynchronous cultures (no delay). Cells delayed in mitosis for a short period showed a similar G1 duration of 9.5-9.7 h ± 3.5 h (Noc^Short^ in Fig. 5a and S8c) to unperturbed cells (AS^No^ ^delay^ Control in Fig. 5b and S8c). However, even a short delay in mitosis increased the proportion of cells arrested in G1 for at least 20 h to 52% (Fig. 5a, Noc^Short^) from 8% (Fig. 5b, AS^NoDelay^ Control). In comparison, a long mitotic delay (Noc^Long^) resulted in over 94% G1 arrest, and the length of G1 in the 6% of cells entering S-phase increased to 13.3 h ± 6.0 h (Fig. 5a and S8c, Noc^Long^). In a key test of our hypothesis, addition of 100 nM MD-224 during the short mitotic delay resulted in over 99% arrest in G1, with correspondingly few S-phase cells after a prolonged time >20 h in G1 (Fig. 5a and S8c, Noc^Short^ +MD-224). Importantly, similar results were obtained using a CENP-E inhibitor to prevent chromosome congression (Fig. S9a-S9d, CENP-E_i_^Short^ ± MD-224), demonstrating that the observed effect was independent of the drug used to cause delay mitosis. Shortening the Noc^Short^ delay by addition of an MPS1 inhibitor to collapse the spindle checkpoint, reduced the percentage of in cells arresting in G1 from 52% to 21%, and reduced G1 length in cells entering S-phase to 8.9 h ± 3.1 h, despite the presence of some chromosome segregation defects (Fig. 5c and Fig. S8c, Noc^Short^ → +MPS1i). In all cases the G1 arrest was robust and maintained for at least 60 h, the maximum time imaged in these experiments, and was not observed in hTERT-RPE1 p53^KO^ FUCCI cells for either the long mitotic arrest (Fig. 5a and S8c, Noc^Long^) or when MD-224 was added during a short mitotic arrest (Fig. 5a and S8c, Noc^Short^ +MD-224). Similar results linking MDM2 levels in mitosis to G1 arrest were also obtained in cells entering mitosis in asynchronous cultures without any cell cycle block. MDM2 destruction triggered by transient MD-224 treatment of mitotic cells in asynchronous culture resulted in an extension of G1 from 9.7 ± 3.5 h to 14.8 ± 2.4 h, and increased the level of G1 arrested cells from 8% to 51% (Fig. 5b, compare Control with MD-224). By contrast, transient inhibition of MDM2 activity in mitosis with Nutlin-3A, an MD-224 related compound which blocks the MDM2-p53 interaction but does not trigger MDM2 destruction, did not result in extended G1 or cell cycle arrest in the following G1 (Fig. 5b, compare Control with Nutlin-3A). Reduction of the level of MDM2 in mitosis and not simply inhibition of its activity is therefore critical for G1 arrest. Importantly, treatment of cells in S-phase and G2 with MD-224 did not trigger arrest in G1 phase of the following cell cycle and all cells entered S-phase, although we did observe a lengthened G1 (Fig. 5d). This is consistent with our data showing rapid recovery of MDM2 following washout of MD-224 in asynchronous cell cultures (Fig. S7c). Taken together, these results show that the proportion of cells undergoing a G1 arrest and length of G1 correlate with the increased length of time in mitosis or reduced level of MDM2.

**Fig. 5.**
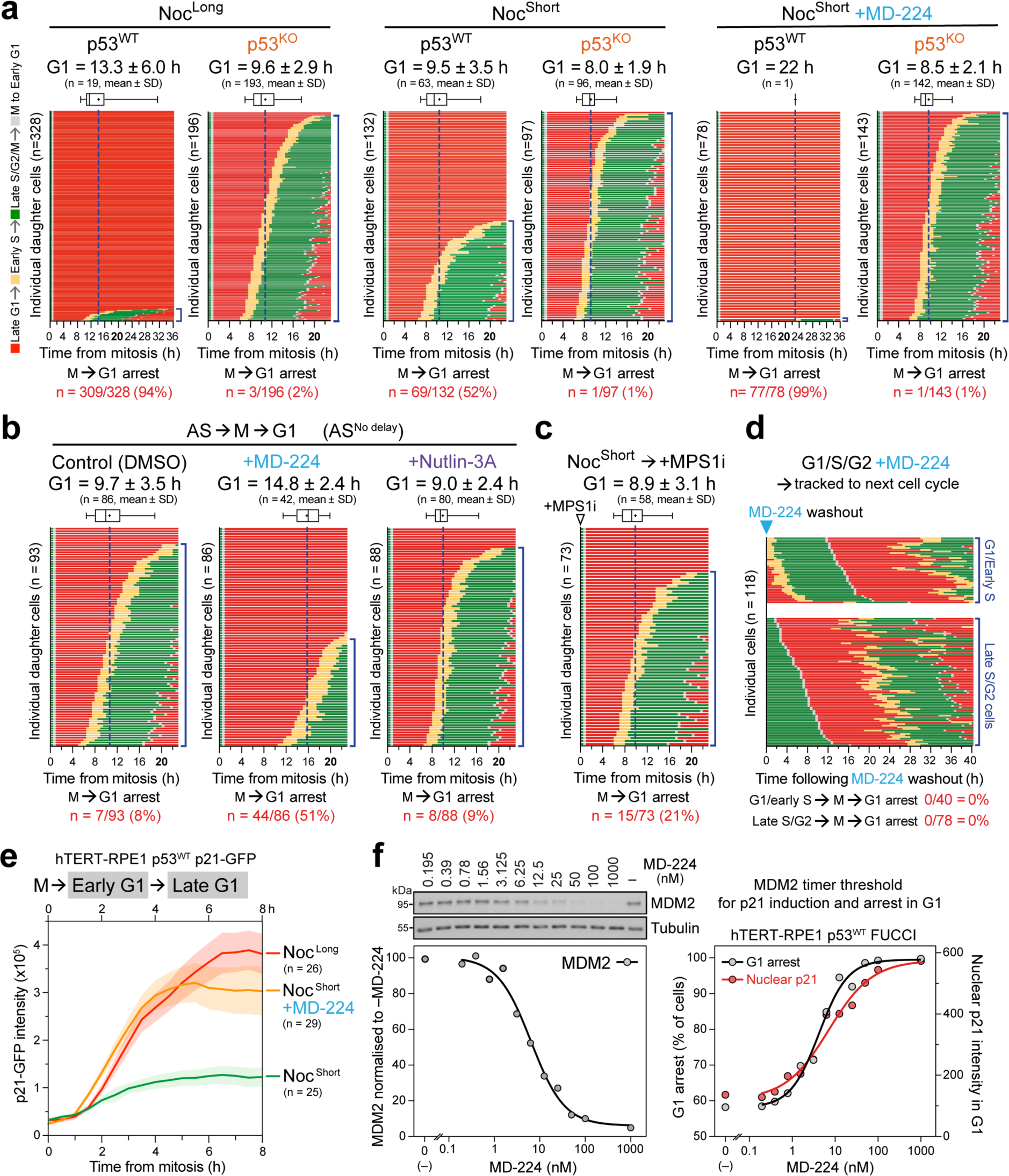
Defining the MDM2 threshold in mitosis for p21 induction and robust cell cycle arrest in G1. **a.** Single cell traces of hTERT-RPE1 p53^WT^ and p53^KO^ FUCCI cells passing from mitosis into the following cell cycle after arrest in mitosis for either 30 min (Noc^Short^) or 4 h (Noc^Long^) with 25 ng/ml nocodazole, in the absence (–) or presence (+) of 100 nM MD-224 for 30 min prior to washout. The mitotic cells were tracked after washout and imaged up to 60 h post-mitosis. The number and proportion of cells arresting in G1 and cells entering S phase are shown. The duration of G1 to cells entering S-phase is shown for all conditions (mean ± SD, sample sizes indicated in the figure). The dotted line in each panel of traces indicates the mean length of G1. A box and whiskers plot show the mean G1 length (+) and 95% confidence intervals. **b.** Mitotic hTERT-RPE1 p53^WT^ FUCCI cells in asynchronous culture (AS^NoDelay^) were treated for 30 minutes with DMSO (Control, n=93), MD-224 (n=86) or Nutlin-3A (n=88), the drugs washed out, and cells then tracked into the following cell cycle. **c.** MPS1 inhibitor was used to override the Noc^Short^ spindle assembly checkpoint arrest. Single cell traces of hTERT-RPE1 p53^WT^ FUCCI cells in the Noc^Short^ + MPS1i condition passing from mitosis into the following cell cycle. The duration of G1 in cells entering S-phase is shown (mean ± SD, n=73). The dotted line in each panel of traces indicates the mean length of G1. The number and proportion of cells arresting in G1 and cells entering S-phase are indicated. **d.** Single cell traces of hTERT-RPE1 p53^WT^ FUCCI cells in either G1/S or S/G2 in asynchronous cultures were treated with MD-224 for 30 min as in the Noc^Short^ condition, washed to remove the drug, then tracked into the following cell cycle. The proportion of cells arresting in G1 and entering S-phase are shown (mean ± SD, n=118). **e.** Levels of p21 in single hTERT-RPE1 p53^WT^ p21-GFP cells were followed post mitosis in new G1 cells for the Noc^Long^ (n=26), Noc^Short^ (n=25) and Noc^Short^+MD-224 (n=29) conditions (mean ± SEM plotted on the graph). **f.** Titration of MD-224 to reveal the threshold concentration required for G1 arrest (black line) of hTERT-RPE1 p53^WT^ FUCCI cells. MDM2 levels were determined by Western blotting (mean, n=2), and p21 levels (mean, red) and proportion of cells arrested in G1 (black) by immunofluorescence microscopy (n=1238-4723 cells for each MD-224 concentration).

To understand the consequences of delays in mitosis and accompanying reduced levels of MDM2 for early events in the following G1, we then measured p21 levels in single hTERT-RPE1 p53^WT^ p21-GFP cells. We observed that after a short 1 h mitotic arrest the level of p21 was slightly increased in early G1 cells, whereas after a longer 4 h delay, or when 100 nM MD-224 was added during the short mitotic delay and then removed, p21 levels rose sharply in early G1, plateaued and were then maintained into late G1 (Fig. 5e). To determine the precise threshold below which MDM2 has to be reduced in mitosis to trigger p21 induction in G1, we titrated MD-224 and measured both the amount of MDM2 by Western blotting, and cell cycle arrest using the hTERT-RPE1 p53^WT^ FUCCI cell line. This revealed a sharp dose-response relationship, where G1 arrest is closely correlated with the increasing level of p21 and decreasing amount of MDM2 (Fig. 5f). The 100 nM concentration of MD-224 used for other experiments sits at the top of this sigmoidal curve, explaining its potency. Taken together, our data therefore reveal a defined threshold for MDM2 in mitosis, below which p53 stabilisation and hence p21 induction in G1 cells leads to cell cycle arrest.

### MDM2 overexpression increases the time in mitosis threshold for G1 arrest

We next asked if increasing the level of MDM2 by overexpression can overcome the mitotic timer response by using hTERT-RPE1 p53^WT^ cells with a stably integrated GFP-MDM2 construct (GFP-MDM2^OE^). Western blotting showed that GFP-MDM2^OE^ cells express around 5-fold more GFP-MDM2 than endogenous MDM2 in hTERT-RPE1 p53^WT^ cells (Fig. 6a and 6b; Fig. S10a). Both proteins have a similar half-life of 21-23 minutes (Fig. S10b), however, due to its increased level in the GFP-MDM2^OE^ cells, MDM2 will therefore take longer to drop below the critical threshold for p21 induction. Supporting this idea, the threshold for p21 induction, determined by MD-224 titration, was increased from 8-10 nM in hTERT-RPE1 p53^WT^ to 25-27 nM MD-224 in GFP-MDM2^OE^ cells (Fig. 6b and S10c). The response of these cells to 1 or 4 h mitotic delay was then compared in mitotic timer and cell proliferation assays. Compared to control hTERT-RPE1 p53^WT^ cells, GFP-MDM2^OE^ cells failed to stabilise p53 or induce p21 in G1 following a 4 h delay in mitosis (Fig. 6c and 6d). Furthermore, GFP-MDM2^OE^ cells did not exhibit the characteristic arrest of cell proliferation observed for the hTERT-RPE1 p53^WT^ cells after a 4-h delay in mitosis (Fig. 6e). Increasing the steady-state level of MDM2 therefore suppresses the mitotic timer response, lending further support to the idea that MDM2 concentration and hence p53 stabilisation is the crucial determinant reporting when the length of mitosis exceeds a crucial time threshold.

**Fig. 6.**
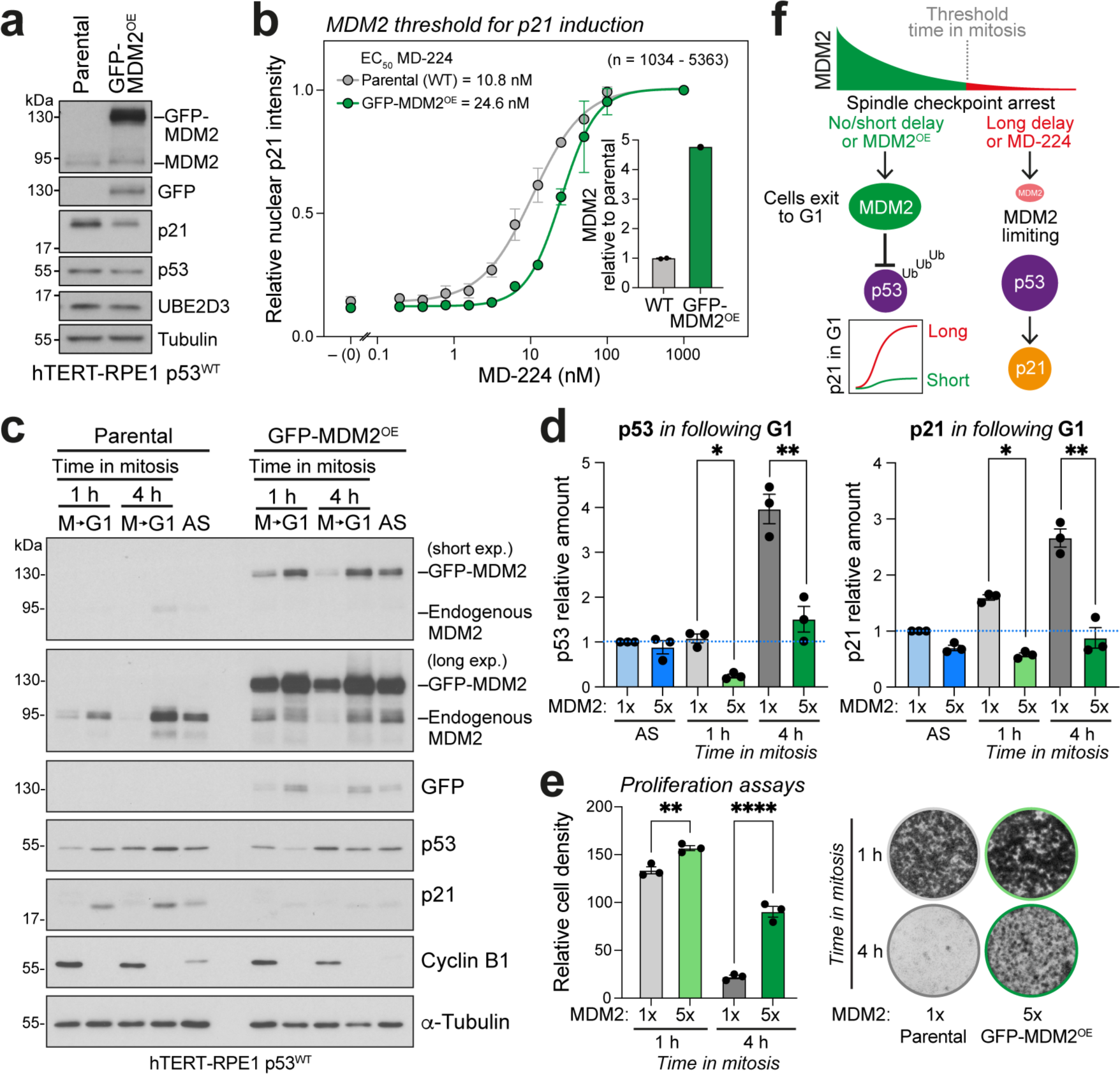
MDM2 overexpression attenuates the mitotic timer response. **a.** Western blot of MDM2 levels in hTERT-RPE1 p53^WT^ (parental) and GFP-MDM2^OE^ cells. **b.** Titration of MD-224 to reveal the threshold concentration required for p21 induction. Nuclear p21 levels were measured by immunofluorescence microscopy in hTERT-RPE1 p53^WT^ (grey line, circles) and GFP-MDM2^OE^ cells (green line, circles). The relative level of MDM2 in GFP-MDM2^OE^ cells compared to parental cells is shown in the bar graph calculated from Figure S10a. **c.** Western blot of cyclin B1 positive mitotic (M) and cyclin B1 negative (G1) hTERT-RPE1 p53^WT^ (parental) or GFP-MDM2^OE^ cells arrested for either 1 h or 4 h in mitosis with 25 ng/ml nocodazole. **d.** Graphs showing the relationship between mitotic delay and the amount of p53 and p21 in the following G1 in parental and GFP-MDM2^OE^ hTERT-RPE1 p53^WT^ cells (mean ± SEM, n=3). Asynchronous hTERT-RPE1 p53^WT^ cells (AS) were used as a normalisation control for the graphs to allow direct comparison of the steady-state levels of p53 and p21. **e.** hTERT-RPE1 p53^WT^ (parental) or GFP-MDM2^OE^ cells were arrested in mitosis for 1 or 4 h with 25 ng/ml nocodazole. Mitotic cells were harvested, washed-out from nocodazole, and 5000 cells plated per well. After 5 days cell proliferation was measured by crystal violet staining (mean ± SEM, n=3). Representative stained wells are shown. **f.** A schematic depicting the decay in MDM2 levels during mitosis. If MDM2 concentration drops below a threshold in mitosis, MDM2 becomes limiting for p53 regulation in the ensuing G1, leading to p53 stabilisation, p21 induction and G1 cell cycle arrest.

## Discussion

These findings support the conclusion that MDM2, due to its short half-life, is a key timer component in the mechanism that triggers a robust cell cycle arrest in the G1 following a prolonged delay in mitosis ^21,22^. MDM2’s timer properties arise through a self-catalysed ubiquitination and proteasomal destruction mechanism, and the attenuation of protein synthesis in mitosis. When MDM2 drops below a threshold concentration in mitosis, it becomes limiting for p53 destruction, leading to p53 stabilisation and p21 induction in new G1 cells (Fig. 6f). Other studies suggest that a PLK1-regulated complex of p53BP1 and USP28, the MDM2 counteracting p53-deubiquitinating enzyme, is also important for G1 arrest following delayed mitosis ^7–1031^. This complex shows the opposite behaviour to MDM2, increasing during mitotic delays, which given the role of USP28 as an antagonist of MDM2, would further stabilise p53 in new G1 cells. However, as G1 arrest behaviour can be uncoupled from time in mitosis using MD-224, we conclude that MDM2 concentration is the key limiting component in the mitotic timer pathway. Our observations help provide an explanation for p53-dependent, yet DNA damage signalling-independent cell cycle arrest in G1 following delays in mitosis ^1–10^. Loss of function mutations or deletion of p53 are amongst the most common genetic changes associated with aneuploid cancers, and we propose this is in part due to the central role of MDM2 in the mitotic timer pathway. Highly aneuploid tumour cells often show prolonged mitosis due to extended spindle assembly checkpoint activation caused by mitotic defects ^1–10^. As long as wild-type p53 functionality is present, the mechanism we have described here will channel these cells into a prolonged cell cycle arrest in the following G1. By contrast, loss of this protective response enables aneuploid cells or cells harbouring other changes that delay progression through mitosis to evade G1 arrest and continue proliferating. These findings shed new light on to the functions of MDM2 and have potentially important consequences for our understanding of how aneuploidy is detected in normal cells, and how failure of this process facilitates genome instability and unchecked proliferation in cancers.

## Acknowledgments

We thank Prof. Ulrike Grüneberg, Prof. Lars Jansen, Prof. Bela Novak, Dr. Madhusudhan Srinivasan, and our colleagues in the Barr group for discussion and advice, Dr. Martin Attwood for help with generation and validation of the p53^KO^ hTERT-RPE1 cells, and Dr. Pawel Mikulski for assistance with RT-qPCR.

## Funding

Cancer Research UK program grant award C20079/A24743 (FAB), and Cancer Research UK Career Development Fellowship C62538/A24670 (IGS).

## Author contributions

Conceptualization: FAB; Methodology: LJF, TS, IGS, FAB; Investigation: LJF, TS, CB; Funding acquisition: FAB, IGS; Supervision: FAB; Writing – original draft: FAB; Writing – review and editing: FAB, LJF, TS.

## Competing interests

The authors declare that they have no competing interests.

## Data and materials availability

All data are available in the main text or the supplementary materials. Reagents generated in this study can be obtained by writing to the corresponding author.

---

**Fig. S1.**
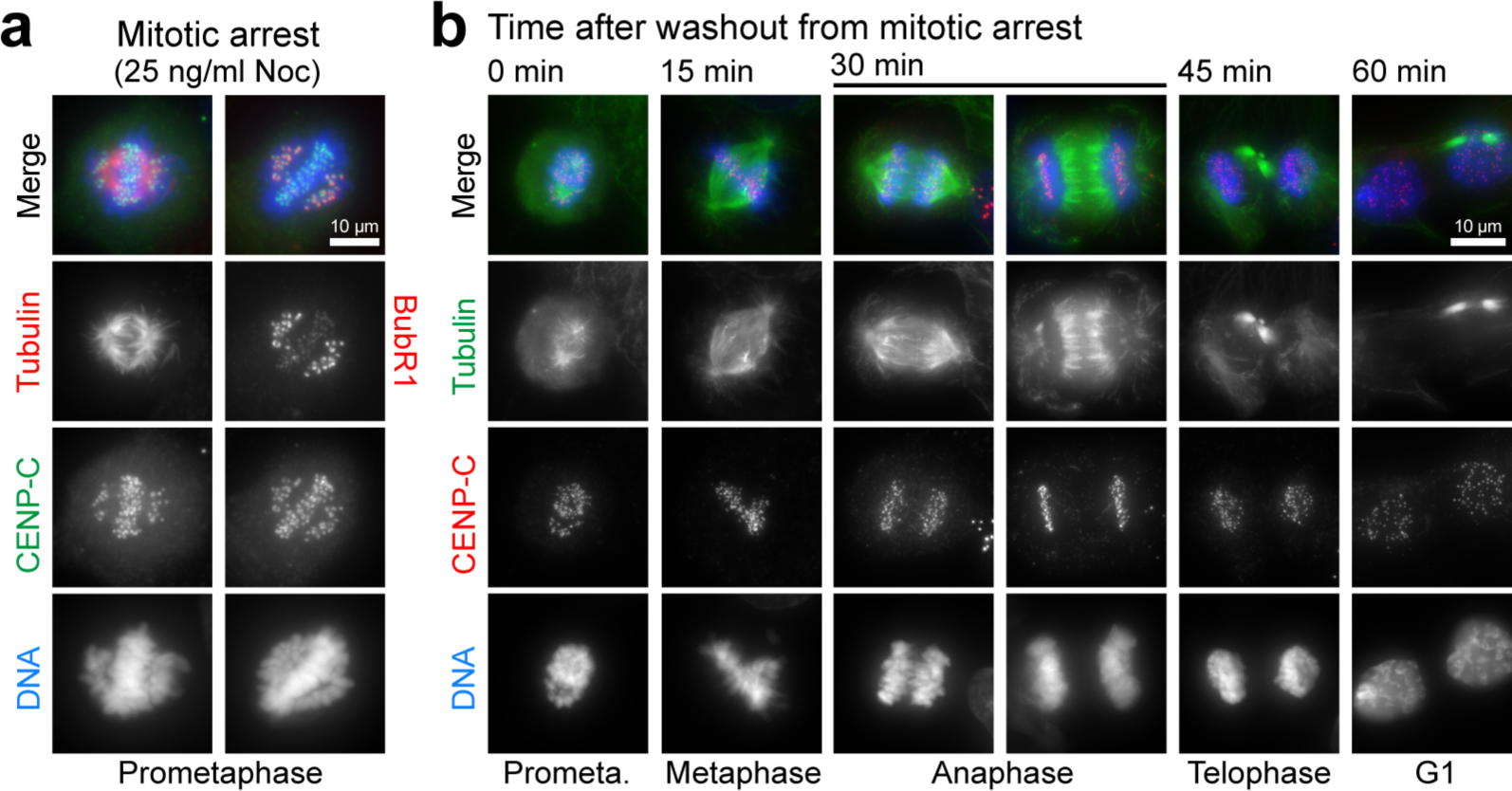
Low dose nocodazole activates the spindle assembly checkpoint and allows rapid release into anaphase. **a.** hTERT-RPE1 cells treated with 25 ng/ml nocodazole (Noc) for 3 h were fixed and stained for DNA, tubulin, the centromere protein CENP-C, and either tubulin or the spindle assembly checkpoint marker BubR1. **b.** Nocodazole was removed by washing, and cells were imaged at the specified time points as they exited mitosis and entered G1.

**Fig. S2.**
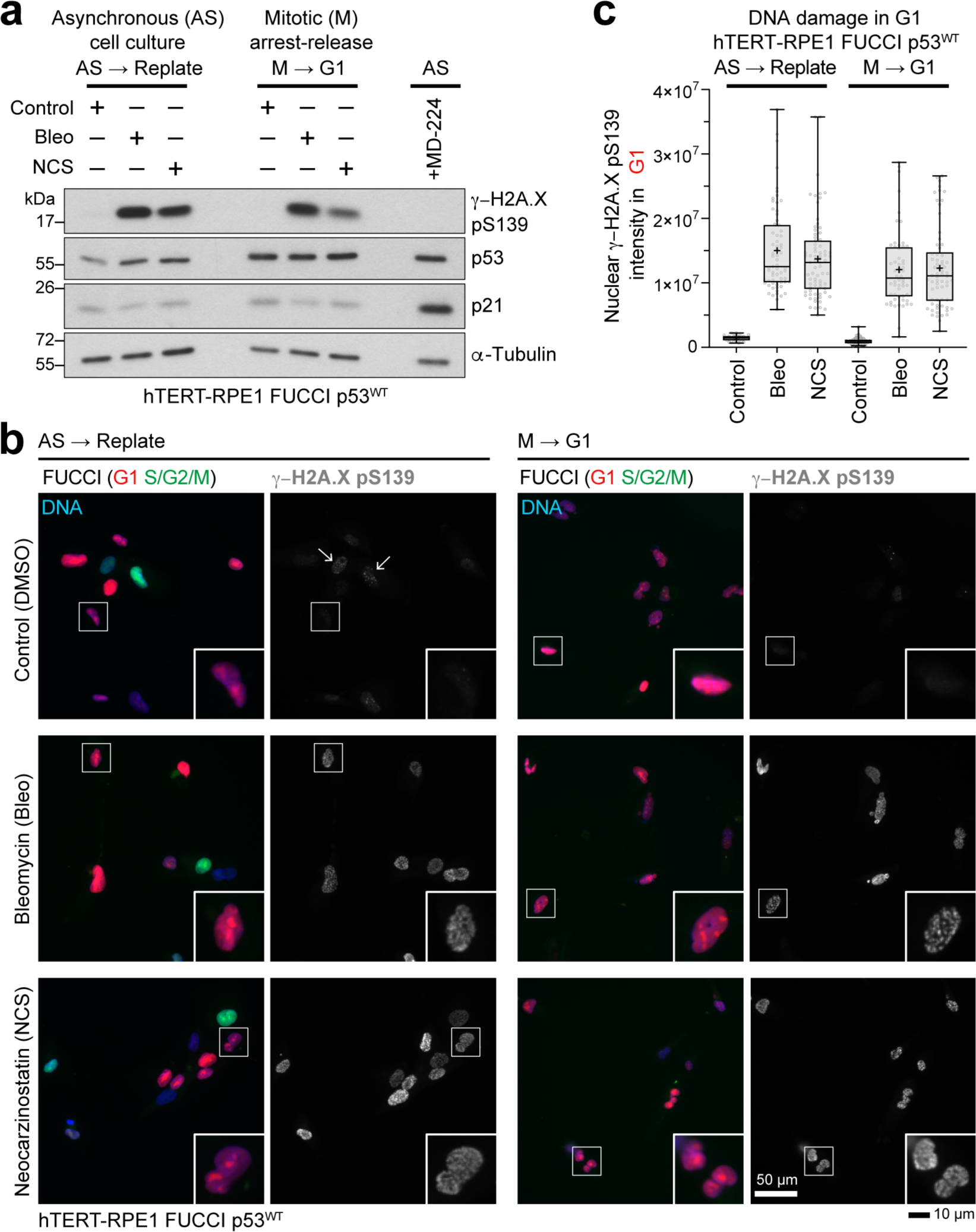
Delays in mitosis do not induce DNA damage in subsequent G1 cells. **a.** Asynchronous (AS) hTERT-RPE1 p53^WT^ cell cultures were plated to mimic the conditions in Fig. 1a-1c, then 17 h later the replated cells were treated with DMSO (Control), 30 µg/ml bleomycin (Bleo) or 200 ng/ml neocarzinostatin (NCS) for 1 h to induce DNA damage. To test the effect of mitotic arrest on DNA damage, hTERT-RPE1 p53^WT^ cells were treated with 25 ng/ml nocodazole for 18 h. Mitotic cells harvested by shake-off and nocodazole washed-out, replated, and then 17 h later treated with DMSO, Bleo or NCS for 1 h. In both cases, DNA damage induction was assessed by Western blotting. **b.** hTERT-RPE1 p53^WT^ FUCCI cells treated as in panel a were stained for γ-H2A.X pS139. The FUCCI marker was directly visualised using fluorescence microscopy. Example G1 cells (red) are shown in the enlarged panels, and arrows indicate S/G2 cells (green) in the AS control condition. Scale bars mark 50 µm, or 10 µm for the enlarged areas marked by white boxes. **c.** DNA damage Box and whiskers plots of nuclear γ-H2A.X pS139 in G1 cells showing the mean (+) for the different conditions.

**Fig. S3.**
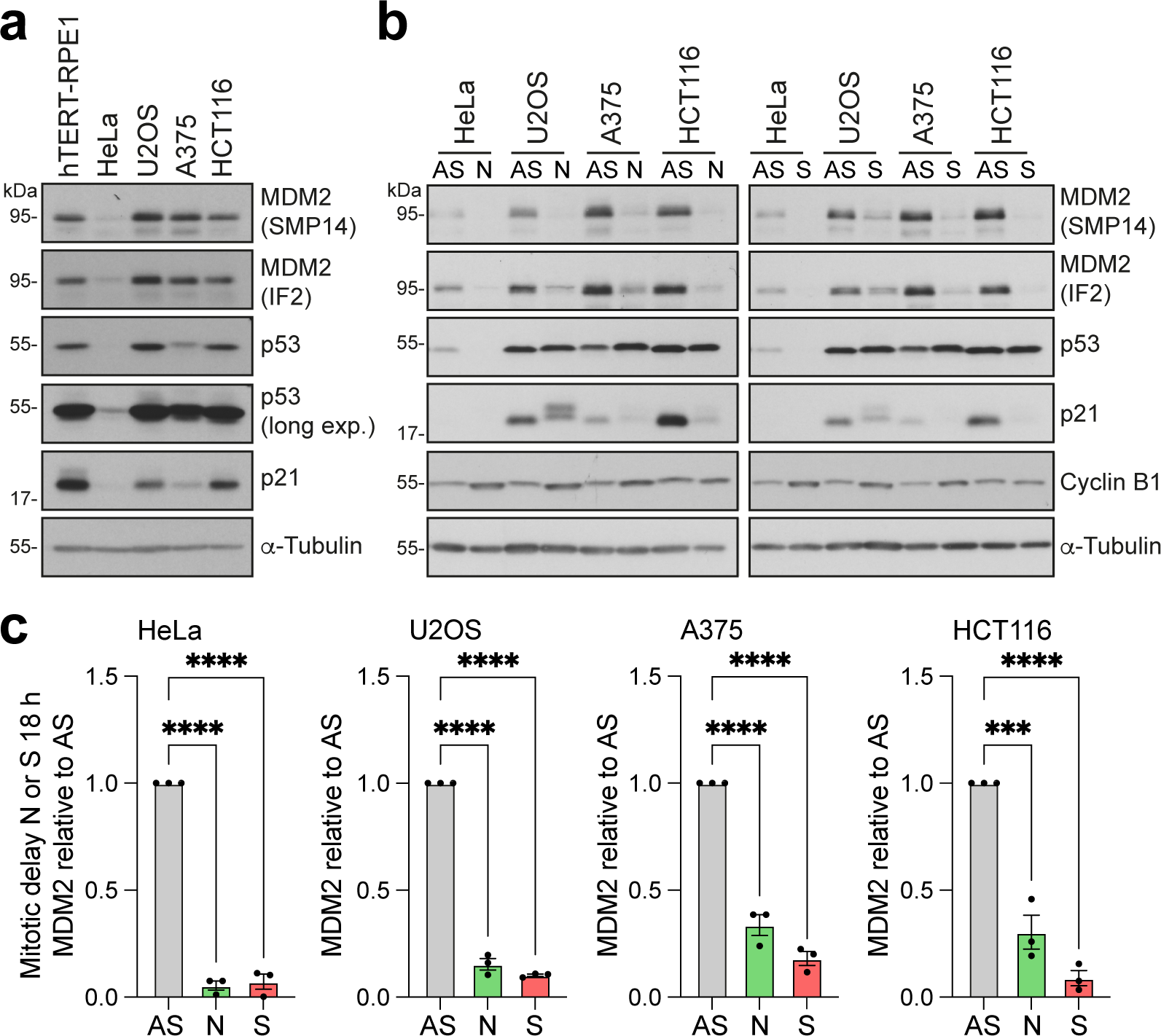
Reduction in the amount of MDM2 during mitosis in different human cell lines. **a.** The amounts of MDM2, p53 and p21 were measured in hTERT-RPE1 p53^WT^, HeLa, U2OS, A375 and HCT116 cell lines under standard culture conditions using Western blotting. α-Tubulin is a loading control. **b.** The amount of MDM2 was measured by Western blotting in asynchronous (AS), nocodazole (N) or S-trityl L-cysteine (STLC, S)-arrested mitotic HeLa, U2OS, A375 and HCT116 cell lines. **c.** Quantification of MDM2 in HeLa, U2OS, A375 and HCT116 cell lines using data from b. Values are expressed relative to the AS control for that cell line (mean ± SEM, n=3).

**Fig. S4.**
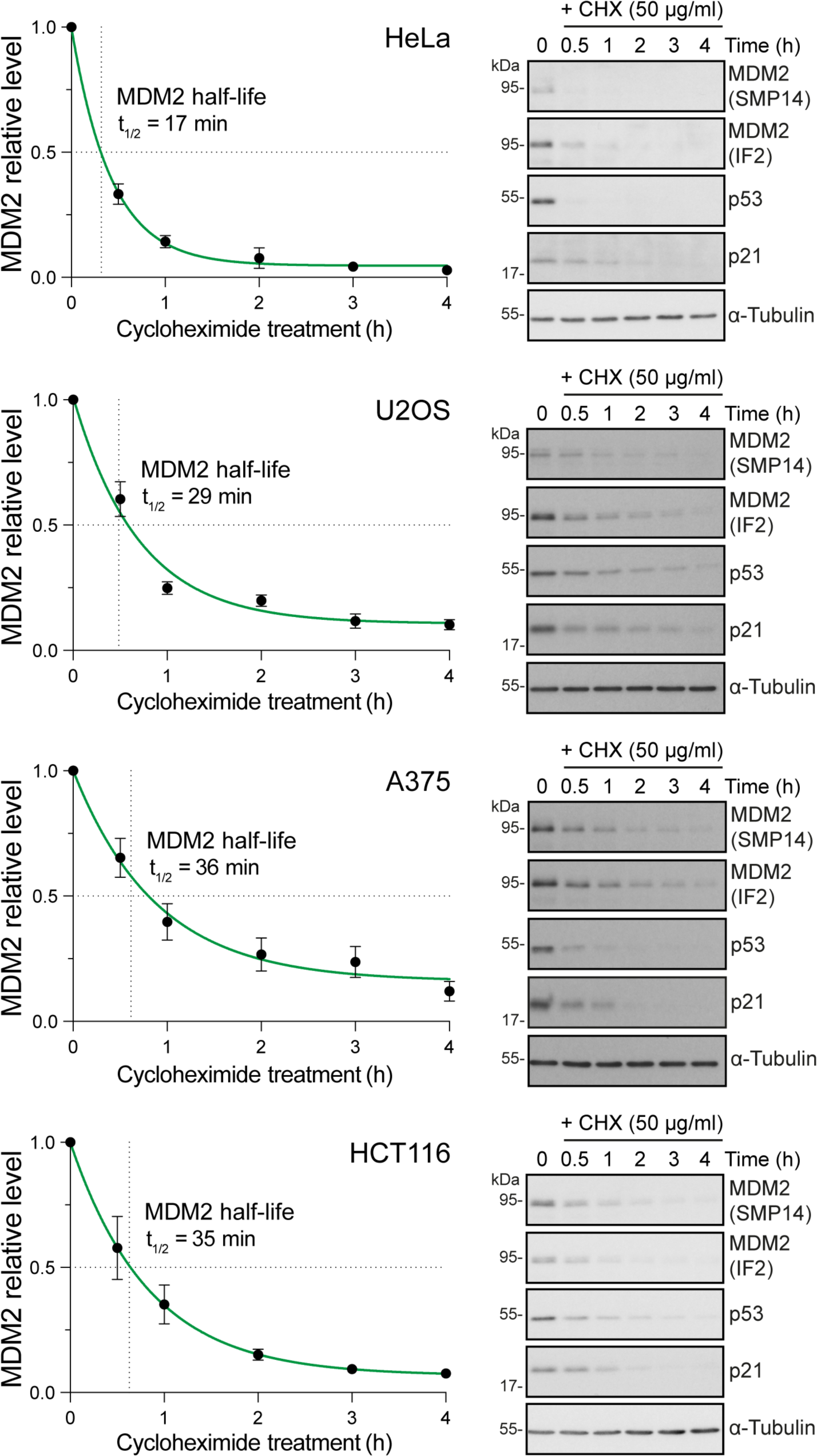
Short half-life is a general property of MDM2 in multiple human cell lines. MDM2 half-life was measured in HeLa, U2OS, A375 and HCT116 cell lines by adding 50 µg/ml cycloheximide (CHX) to block protein synthesis, and then collecting samples up to 4 h. MDM2 half-life is indicated in the graph (mean ± SEM, n=3), with example western blots.

**Fig. S5.**
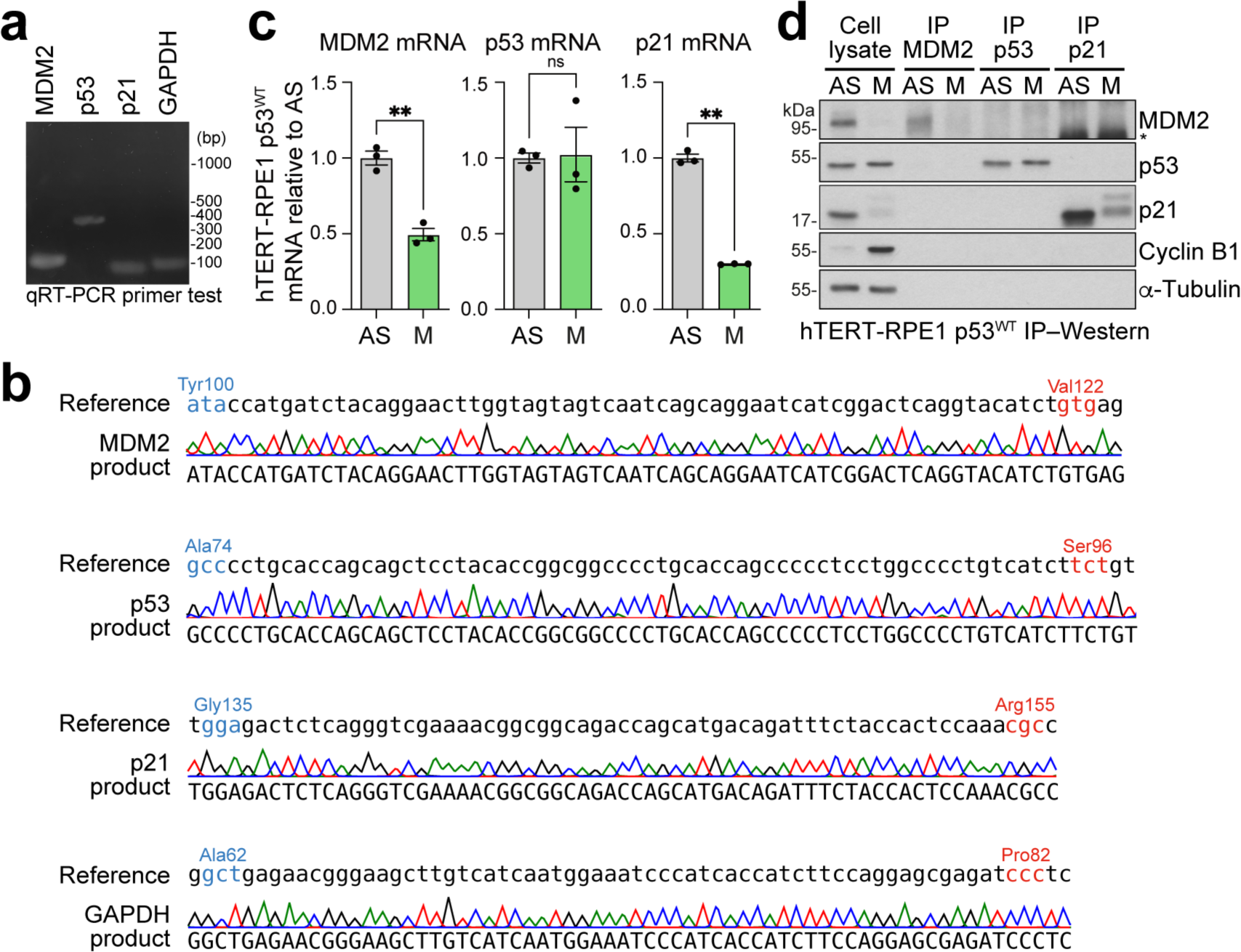
RT-qPCR reveals reduction of MDM2 mRNA in mitosis. **a.** Agarose gel showing RT-qPCR products obtained using diagnostic primers for MDM2, p53, p21 and GAPDH. **b.** Reference and confirmed DNA sequences for the MDM2, p53, p21 and GAPDH RT-qPCR products. **c.** RT-qPCR from asynchronous hTERT-RPE1 p53^WT^ cells (AS) and cells arrested for 18 h in mitosis with 25 ng/ml nocodazole (M). All mRNA levels are expressed relative to the AS control samples, and normalised against GAPDH. **d.** Immunoprecipitation of MDM2, p53 and p21 from AS and M conditions used in panel c. The asterisk (*) denotes a non-specific band.

**Fig. S6.**
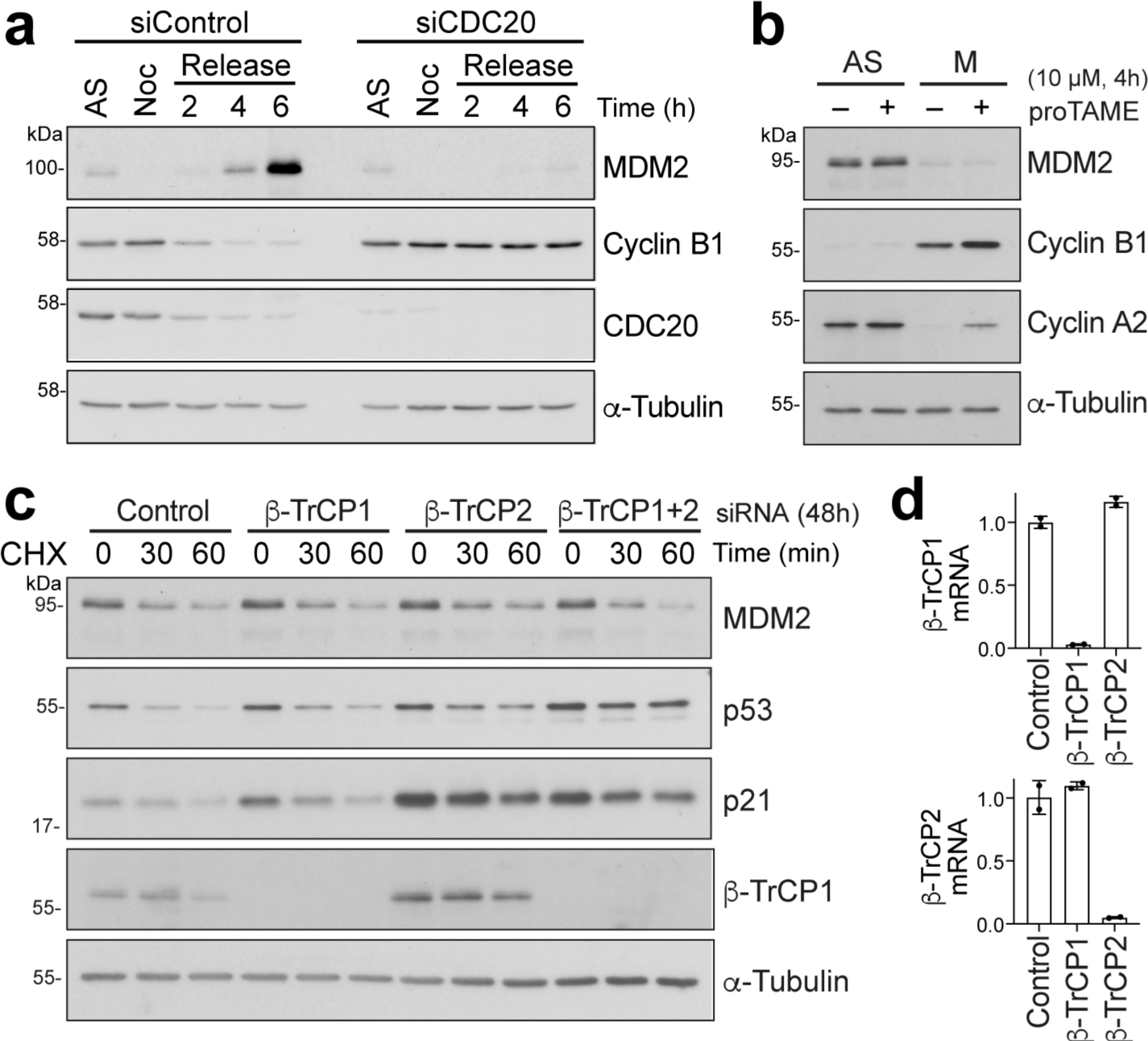
APC/C^CDC20^ and SCF^β-TRCP^ do not regulate MDM2 stability in mitosis. **a.** HeLa cells depleted of the APC/C co-activator CDC20 (siCDC20) or treated with a non-targeting control (siControl) were arrested in mitosis for 18 h with 100 ng/ml nocodazole (Noc). Nocodazole was washed out to test for CDC20 depletion, confirmed by cyclin B1 stabilisation for up to 6 h in the siCDC20 treated cells. Untreated asynchronous cells (AS) were taken as a control. Compared to asynchronous (AS) cells, MDM2 was lost during the mitotic arrest independently of CDC20, and did not recover in the washout in CDC20-depleted cells due to the sustained mitotic arrest. MDM2 recovered in the control cells due to resumption of protein synthesis during mitotic exit and G1 entry. **b.** MDM2 levels were measured in AS and 25 ng/ml nocodazole-arrested mitotic cells by Western blot after 4 h treatment with 10 µM of the APC/C inhibitor proTAME (+). Solvent treated (–) cells were used as a control. **c.** MDM2 stability was measured by Western blot following 30- or 60-minutes treatment with 50 µg/ml cycloheximide (CHX) in cells depleted of the SCF subunits β-TRCP1, β-TRCP2, or β-TRCP1 and 2 together. Non-targeting siRNA was used as a control. **d.** RT-qPCR was used to measure the levels of β-TRCP1 and β-TRCP2 mRNAs in cells depleted of β-TrCP1 or β-TrCP2 by siRNA to confirm successful knockdown. Signals were normalised to GAPDH mRNA and expressed relative to the Control sample, and the mean ± SD (n=2) is plotted in the bar graphs.

**Fig. S7.**
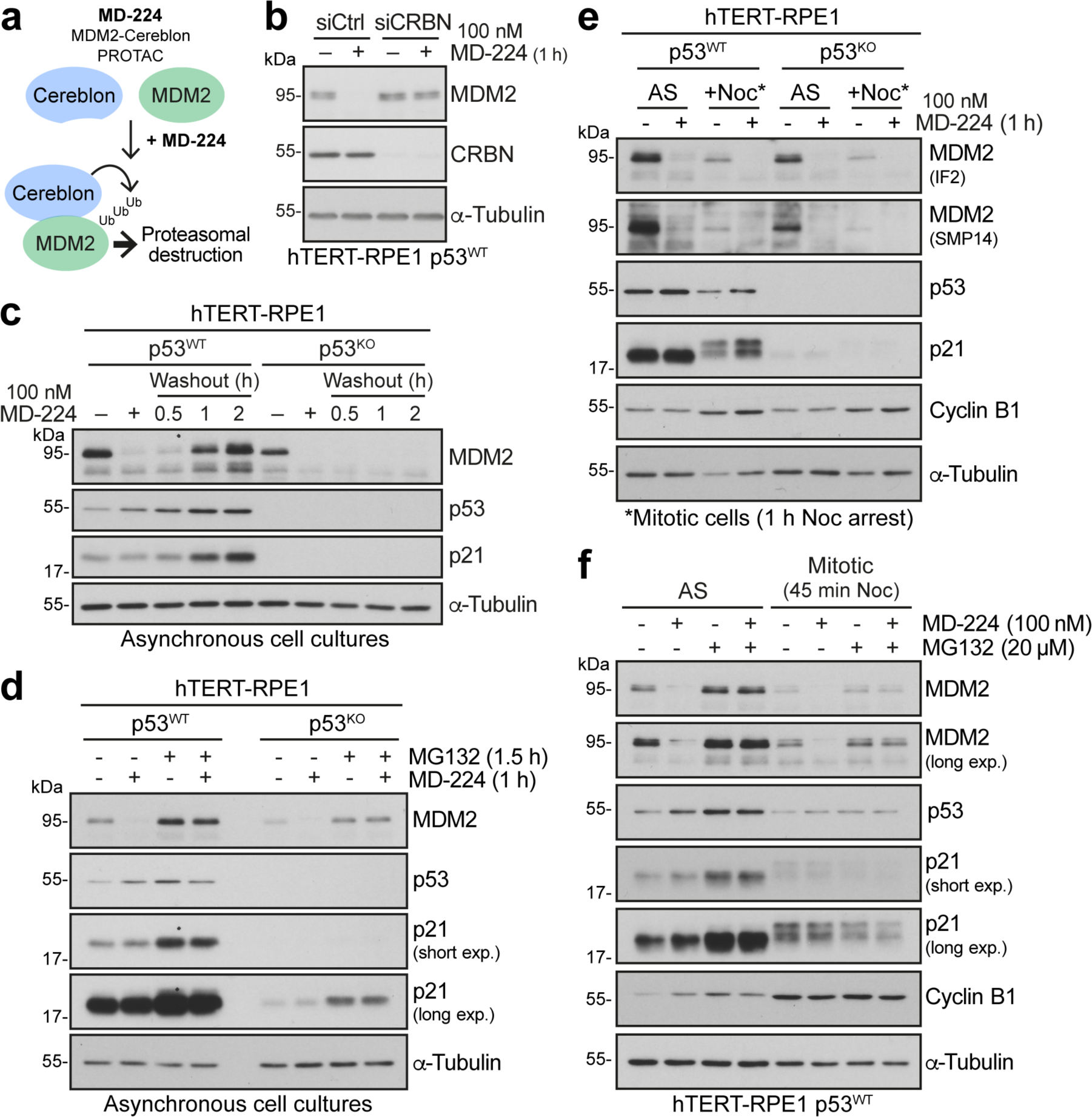
MD-224 results in rapid and reversible MDM2 destruction in hTERT-RPE1 cells. **a.** A cartoon showing the mode of action for MD-224. **b.** hTERT-RPE1 p53^WT^ cells depleted of CRBN with siRNA were treated with solvent (−) or 100 nM MD-224 (+) for 1 h, and Western blotted. MD-224 requires CRBN for MDM2 destruction. Non-targeting siRNA was used as a control. **c.** hTERT-RPE1 p53^WT^ or p53^KO^ cells were treated with 100 nM MD-224 for 1 h (+) or a solvent control (–). MD-224 was then washed out with fresh growth medium for 0.5 to 2 h and Western blotted for MDM2, p53 and p21. Tubulin was used as a loading control. **d.** The amount of MDM2 was measured by Western blotting in asynchronous (AS) hTERT-RPE1 p53^WT^ and p53^KO^ cells treated for 1.5 h with (+) or without (–) the proteasome inhibitor MG132 (20 µM) and for 1 h with (+) or without (–) 100 nM MD-224. **e.** The amount of MDM2 was measured in asynchronous cells (AS) or mitotic hTERT-RPE1 p53^WT^ and p53^KO^ cells arrested for 1 h with 25 ng/ml nocodazole (+Noc) in the presence (+) or absence (–) of 100 nM MD-224. MDM2 was detected by Western blot using two different antibodies. **f.** The amount of MDM2 was measured using Western blotting in asynchronous cells (AS) or mitotic hTERT-RPE1 p53^WT^ cells arrested for 45 min with 25 ng/ml nocodazole (+Noc) in the presence (+) or absence (–) of 100 nM MD-224 and the proteasome inhibitor MG132 (20 µM).

**Fig. S8.**
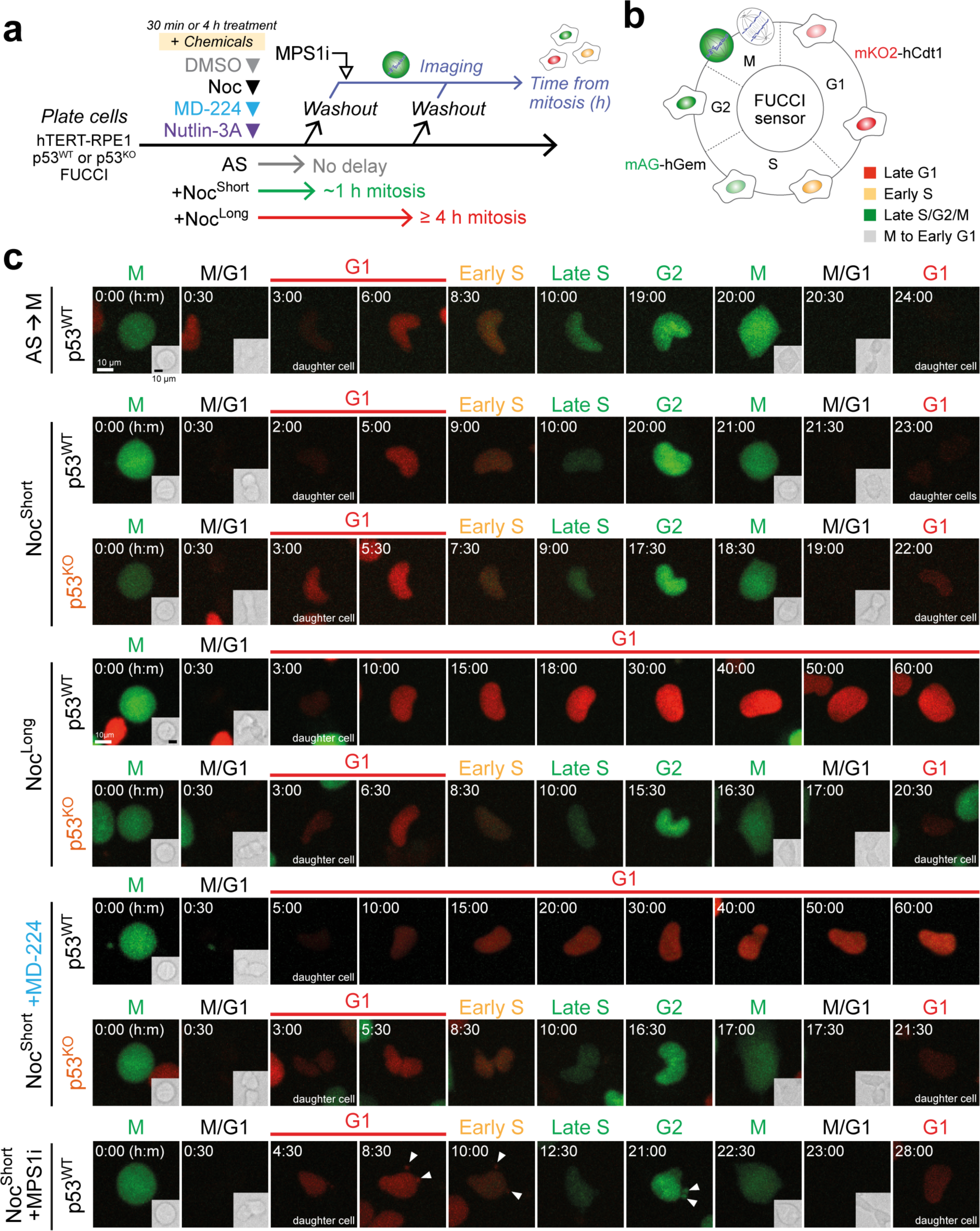
hTERT-RPE1 FUCCI cell lines progress through G1 into S-phase when mitosis is not delayed. **a.** Experimental design used for Figure 5. **b.** The properties of the FUCCI sensor. **c.** Representative images of hTERT-RPE1 p53^WT^ and p53^KO^ FUCCI cells for the conditions described in Figure 5. Arrowheads in Noc^Short^+MPS1i indicate micronuclei.

**Fig. S9.**
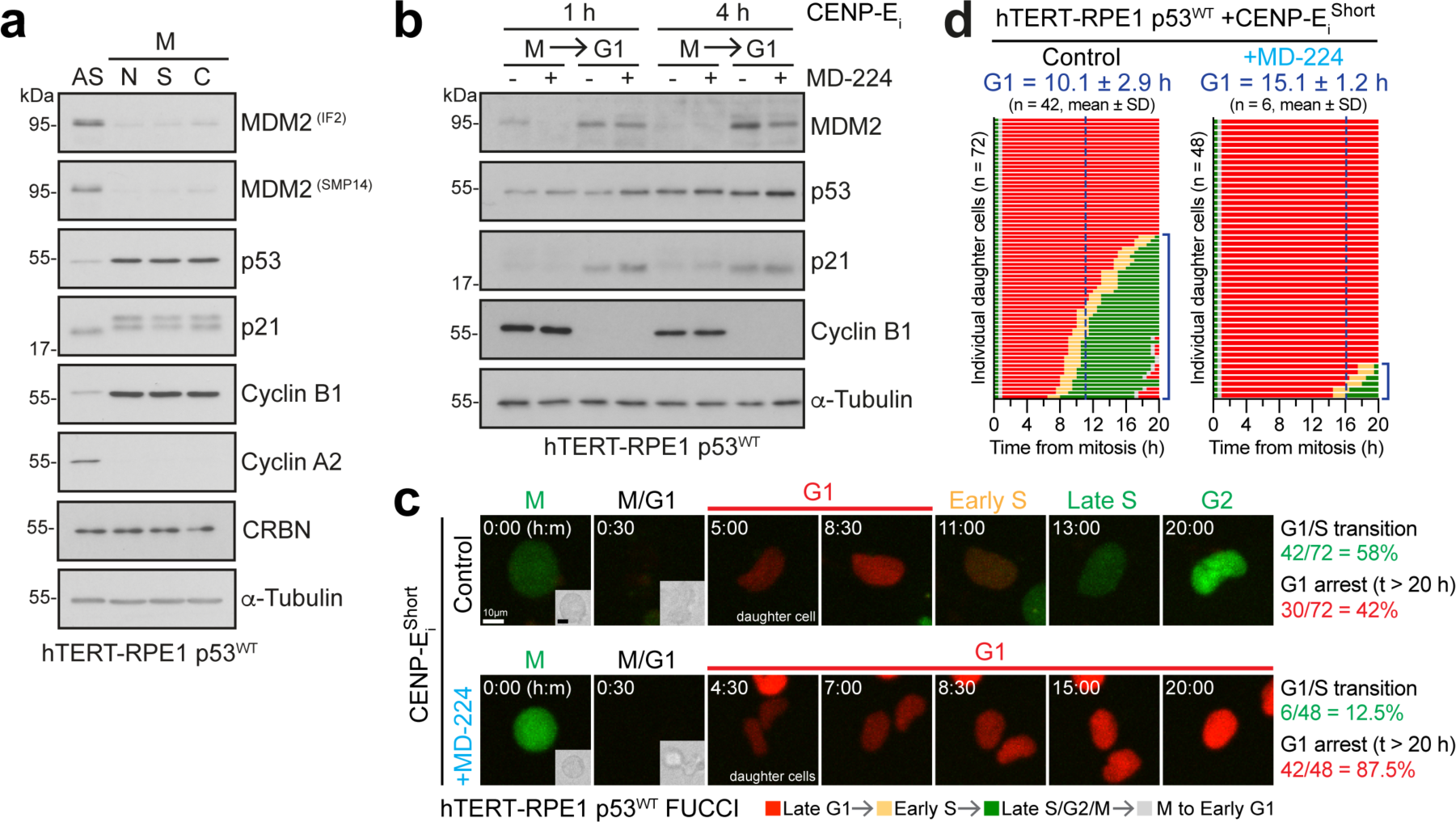
CENP-E inhibitor figure. **a.** The amount of MDM2 was measured by Western blotting in asynchronous (AS) or mitotic hTERT-RPE1 p53^WT^ cells, arrested in mitosis for 18 h with 100 ng/ml nocodazole (N), 5 µM S-trityl L-cysteine (S), or 300 nM CENP-E inhibitor (C). **b.** Western blot of cyclin B1 positive mitotic (M) and cyclin B1 negative (G1) hTERT-RPE1 p53^WT^ cells arrested for either 1 h or 4 h in mitosis with 30 nM CENPE-inhibitor (CENPE_i_), in the absence (–) or presence (+) of 100 nM MD-224 for the final hour. **c.** Representative images of hTERT-RPE1 p53^WT^ passing from mitosis into the following cell cycle after arrest in mitosis for 45 min with 30 nM CENP-E inhibitor (CENP-E_i_^Short^) in the absence or presence of 100 nM MD-224 prior to washout. The mitotic cells were tracked after washout and then imaged. The number and proportion of cells arresting in G1 and cells entering S phase are shown. **d.** Single cell traces of hTERT-RPE1 p53^WT^ FUCCI cells treated as in panel e. passing from mitosis into the following cell cycle. The duration of G1 to cells entering S-phase is shown for all conditions (mean ± SD, sample sizes indicated in the figure).

**Fig. S10.**
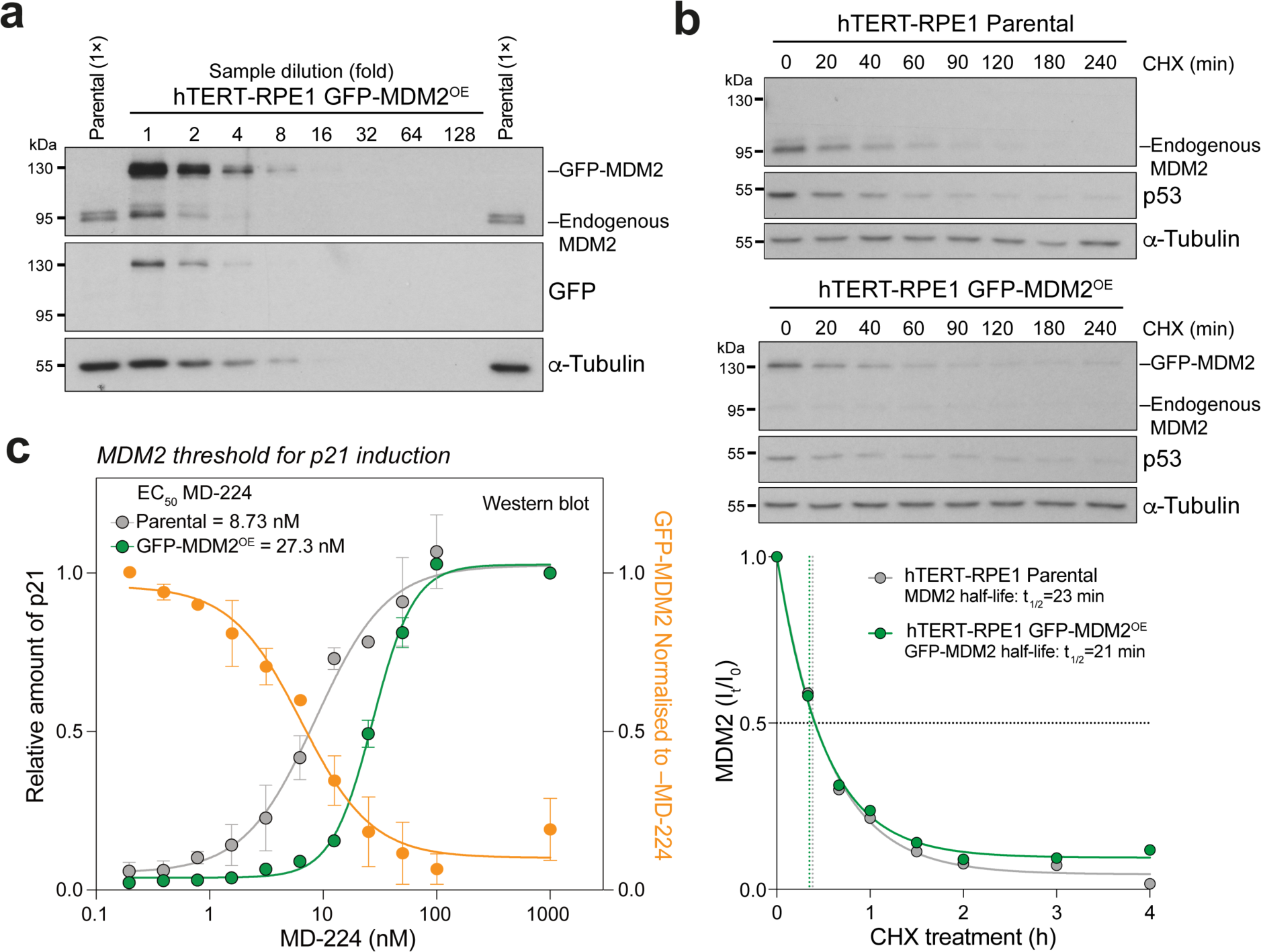
Additional characterisation of hTERT1-RPE1 GFP-MDM2 overexpressing cells. **a.** Western blot of undiluted parental hTERT-RPE1 p53^WT^ and a dilution series of GFP-MDM2^OE^ cell lysates. GFP-MDM2 level relative to endogenous MDM2 was measured from this data. **b.** MDM2 and GFP-MDM2 half-lives were measured using cycloheximide treatment of parental hTERT-RPE1 p53^WT^ and GFP-MDM2^OE^ cells for up to 4 h. Samples were Western blotted and MDM2 levels measured and plotted in the graphs. Half-lives were estimated using a 1-phase curve fit. **c.** Titration of MD-224 to reveal the threshold MDM2 concentration (orange line, circles) required for p21 induction. Total p21 levels were measured by Western blot in hTERT-RPE1 p53^WT^ (grey line, circles) and GFP-MDM2^OE^ cells (green line, circles) (mean ± SEM, n=2). GFP-MDM2 levels in GFP-MDM2^OE^ cells were also measured (orange line, circles) (mean ± SEM, n=2)

## Methods

### Reagents and antibodies

General laboratory chemicals were obtained from Merck, Sigma-Aldrich, and Thermo Fisher Scientific. Commercially available polyclonal antibodies (pAbs) or monoclonal antibodies (mAbs) used for Western blotting (WB), immunofluorescence (IF), and immunoprecipitation (IP), are listed in Table 1.

**Table 1:** Primary and secondary antibodies used in this study.

| Antibody target | Company | mAb/pAb | Catalogue number | Species | Dilution (WB) | Dilution (IF) | Notes |
| --- | --- | --- | --- | --- | --- | --- | --- |
| MDM2 (IF2) | Millipore | mAb | OP46 | Mouse | 1:1000 |  |  |
| MDM2 (SMP14) | Abcam | mAb | Ab3110 | Mouse | 1:1000 |  |  |
| MDM2 (D1V2Z) | CST | mAb | 86934 | Rabbit | 1:1000 |  | Used for IP |
| p53 (DO-7) | CST | mAb | 48818 | Mouse | 1:1000 | 1:1000 |  |
| p53 (7F5) | CST | mAb | 2527 | Rabbit |  |  | Used for IP |
| p21 (12D1) | CST | mAb | 2947 | Rabbit | 1:1000 | 1:1000 | Used for IP |
| p21 (CP36/CP74) | Millipore | mAb | 05-345 | Mouse | 1:1000 |  |  |
| UBE2D2/Ubc H5b | Novus Bio | pAb | NPB1-81769 | Rabbit | 1:1000 |  |  |
| UBE2D3/Ubc H5C (D60E2) | CST | mAb | 4330 | Rabbit | 1:1000 |  |  |
| CENP-C | MBL | pAb | PD030 | Guinea Pig |  | 1:1000 |  |
| BubR1 (8G1) | Millipore | mAb | MAB3612 | Mouse |  | 1:1000 |  |
| Cyclin B1 (GSN3) | Millipore | mAb | 8A5D12/05-373 | Mouse | 1:5000 |  |  |
| Cyclin A2 (E399) | Abcam | mAb | Ab32498 | Rabbit | 1:500 |  |  |
| CRBN (D8H3S) | CST | mAb | 71810 | Rabbit | 1:1000 |  |  |
| α-Tubulin (DM1A) | Sigma | mAb | T6199 | Mouse | 1:10000 | 1:1000 | |
| Cdc20 | Proteintech | pAb | 10252-1-AP | Rabbit | 1:1000 |  |  |
| β-TrCP1 (D13F10) | CST | mAb | 4394 | Rabbit | 1:1000 | | |
| GFP (7.1 and 13.1) | Roche | mAb | 11814460001 | Mouse | 1:1000 |  |  |
| yH2A.X<br>pS139 (2F3) | BioLegend | mAb | 613401 | Mouse | 1:1000 | 1:1000 |  |
| Mouse-HRP | Jackson | pAb | 715-035-150 | Donkey | 1:5000 |  |  |
| Rabbit-HRP | Jackson | pAb | 711-035-152 | Donkey | 1:5000 |  |  |
| Anti-Rabbit<br>AlexaFluor®-<br>488/555/568<br>/647 | Thermo<br>Fisher |  | A21206/A<br>31572/A1<br>0042/A31<br>573 | Donkey |  | 1:1000 |  |
| Anti-Mouse<br>AlexaFluor®-<br>488/555/568<br>/647 | Thermo<br>Fisher |  | A21202/<br>A31570/A<br>10037/A3<br>1571 | Donkey |  | 1:1000 |  |
| Alexa Fluor®<br>647 Anti-Guinea Pig<br>IgG (H+L) | Jackson |  | 706-605-148 | Donkey |  | 1:1000 |  |

### Inhibitors and drug compounds

Commercially-sourced inhibitors or drug compounds used in this study, and the solvents used to reconstitute them, are listed in Table 2. Experiment-specific timings and concentrations are given in the figure legends.

**Table 2:** Small molecules used in this study.

| Drug/inhibitor | Target | Company | Catalogue number | Concentration | Solvent |
| --- | --- | --- | --- | --- | --- |
| Nocodazole | Microtubules | Sigma-Aldrich | 487928 | 25 or 100 ng/ml | DMSO |
| STLC | KIF11 | Sigma-Aldrich | 164739 | 5 µM | DMSO |
| GSK923295 | CENP-E | TOCRIS | 6837 | 30 – 300 nM | DMSO |
| MD-224 | MDM2 | MedChemExpress | HY-114312 | 100 nM | DMSO |
| Nutlin-3a | MDM2 | Selleck Chem | S8059 | 2.5 µM | DMSO |
| Cycloheximide | Ribosome | Sigma-Aldrich | C7698 | 50 µg/ml | Water |
| MG132 | Proteasome | Sigma-Aldrich | 474790 | 20 µM | DMSO |
| AZ3146 | MPS1 | TOCRIS | 3994 | 2 µM | DMSO |
| ProTAME | APC/C | BostonBiochem | I-440 | 10 µM | DMSO |
| Bleomycin sulfate | DNA | Selleck | S1214 | 30 µg/ml | DMSO |
| Neocarzinostatin | DNA | Sigma-Aldrich | N9162 | 200 ng/ml | 20 mM<br>MES, pH<br>7.4 |

### MDM2 expression plasmid construction

Mammalian eGFP-MDM2 expression plasmids were generated using pEF5/FRT or pcDNA5/FRT plasmids harbouring eGFP followed by a 25 amino acid linker YK(GSSS)_5_RIP 5’ to the multiple cloning site, as the backbone. The full-length MDM2 coding sequence was isolated from Addgene plasmid #16233 (pcDNA3 FRT MDM2) by PCR using the KOD hot start polymerase kit (Merck #71086), using primers that included a 5’ BamHI restriction site, and a 3’ XhoI restriction site. Following BamHI (NEB #R3136)-XhoI (NEB #R0146) digest of the MDM2 PCR product and destination vector, ligations were performed with T4 ligase (NEB #M0202). Ligated plasmids were transformed into competent XL-1 blue cells (Agilent Technologies #200249). I440A and R479A MDM2 mutants were generated by site-directed mutagenesis, using KOD polymerase. MDM2 truncated fragments were generated by PCR using Addgene plasmid #16233 (pcDNA3 FRT MDM2) as a template and cloned into pEF5 FRT eGFP with BamHI-XhoI. Primers used to generate constructs, are listed in Table 3. The mammalian expression eGFP-MDM2 overexpression plasmid was generated by subcloning the eGFP-MDM2 insert into a pMK232 vector (Addgene #72834) following NdeI (NEB #R0111S)-XhoI (NEB #R0146) digest.

**Table 3:**
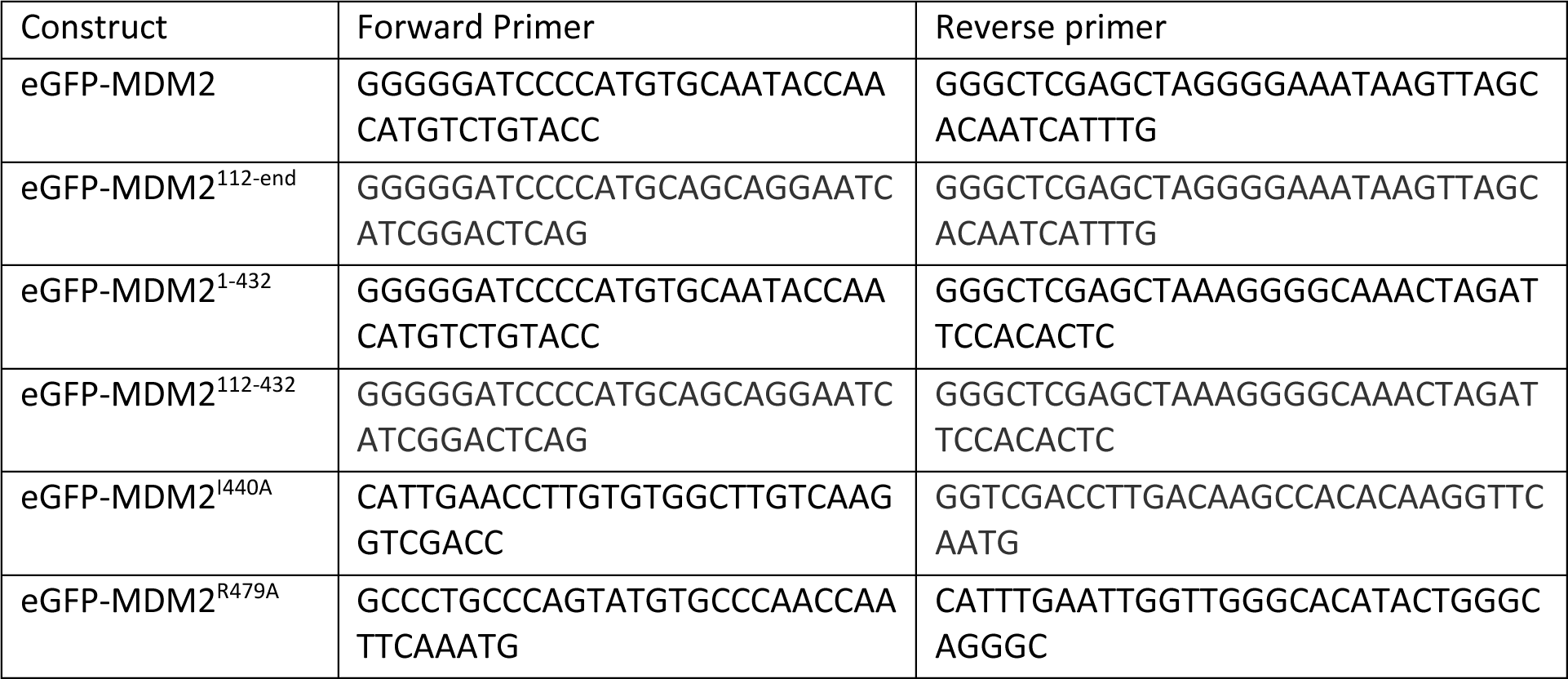
Primers used to generate the MDM2 plasmids used in this study.

### Cell lines and cell culture

All stock cell lines are validated stocks purchased from the ATCC. hTERT-RPE1 cells (#CRL-4000) were cultured in DMEM-F12 Ham medium (Sigma #D6421), supplemented with GlutaMAX™ (Gibco #35050087) and 10% (v/v) fetal bovine serum (FBS, Sigma-Aldrich #F9665). HeLa S3 (#CRL-2.2), A375 (#CRL-1619), U2OS (#HTB-96), and HEK293T (#CRL-3216) cells were cultured in DMEM containing GlutaMAX™ (Gibco #10569010) and 10% (v/v) FBS. HCT116 cells (#CCL-247) were cultured in McCoy’s 5A medium (Gibco #16600082), supplemented with 10 mM sodium pyruvate (Thermo Fisher #11360088) and 10% (v/v) FBS. The hTERT-RPE1 p53^WT^ p21-GFP cell line has been described previously ^32^. For routine passaging of cells, cells were washed in PBS, and incubated with TrypLE™ Express Enzyme cell dissociation reagent (Gibco #12605036) for 5 min at 37°C, before resuspending detached cells in full medium for passage. All cell lines are wild-type for p53. HeLa cells express the HPV E6/E7 proteins which result in p53 downregulation. All cell lines were maintained in a humidified 37°C cell culture incubator (Thermo, HERAcell, #51013568), at 5% CO_2_.

### Transient transfection of cells

Cells were seeded at a density of 50,000 cells per well in 6-well plates. The next day, transfection mixes were prepared in sterile DNase-free Eppendorf tubes (Eppendorf #0030108035) consisting of 100 µl Opti-MEM (Gibco #11058021), 3 µl Mirus TransIT-LT1 transfection reagent (Mirus #MIR2300), and up to 1 µg of plasmid DNA. If less than 1 µg plasmid DNA was required, carrier pBlueScript II (Agilent) empty plasmid was used to bring the total amount of DNA up to 1 µg. Transfection mixes were vortexed briefly for 20 s, and incubated at room temperature for 30 min, before adding drop-wise to cells. Cells were then left to grow for the times indicated in the figure legends. To generate the eGFP-MDM2 overexpressing cell line, 800 ng of the pMK232 eGFP-MDM2 plasmid was transfected into p53^WT^ hTERT-RPE1 cells, as described above. 48 h post-transfection, cells were selected with puromycin and diluted to 1 cell per well of a 96-well plate. Viable colonies were screened for GFP-MDM2 overexpression.

### CRISPR/Cas9 gene editing of hTERT-RPE1 cells

First, hTERT-RPE1 cells were edited to create a functionally null puromycin resistance cassette through delivery of two pX461 vectors (Addgene #48140) containing the appropriate guide RNA (gRNA) sequences (CGCTCAACTCGGCCATGCGC; GCAACAGATGGAAGGCCTCC) using the Amaxa^TM^ nucleofector^TM^ kit V (Lonza #VCA-1003). Monoclonal lines were screened for puromycin sensitivity, growth rate, and cilium formation. We refer to this cell line as hTERT-RPE1 p53^WT^ throughout. Subsequently, to generate p53 knockout cells, the pX459 vector containing guide RNA against p53 (CCATTGTTCAATATCGTCCG) was delivered through electroporation. After brief selection in puromycin, monoclonal lines were screened for Nutlin-3a resistance, followed by extensive quality controls including immunoblotting and amplification of the targeted genomic region by PCR. Genomic DNA was isolated from clones using QuickExtract^TM^ DNA isolation solution (Lucigen #QE09050). PCRs were performed with KOD hot-start polymerase (Merck #71086), using primers flanking the gRNA recognition site. PCR products were ligated into pSC-A vectors using a Blunt-end PCR cloning kit (Agilent #240207). Ligated plasmids were transformed into XL-1 blue competent cells, and around 10 colonies were mini-prepped (Qiagen #27107) and sequenced using Sanger DNA sequencing (Source Genomics, Cambridge, UK) in order to cover all edited alleles. The p53^KO^ cell lines were STR profiled using the ATCC cell line authentication service (ATCC, Manassas, VA), and validated as hTERT-RPE1. We refer to this cell line as hTERT-RPE1 p53^KO^ throughout.

### hTERT-RPE1 FUCCI cell line generation

The hTERT-RPE1 p53^WT^ and p53^KO^ FUCCI cells used here were generated by lentiviral infection. Briefly, pBOB-EF1-FastFUCCI-Puro (Addgene, #86849), pMD2.G (Addgene #12259) and psPAX2 (Addgene #12260) (gifts from Didier Trono) were used for lentiviral-based packaging in HEK293T cells for two days with a media change after 24 h. The resulting lentiviral supernatant was used to infect hTERT-RPE1 p53^WT^ and p53^KO^ for 24 h. After infection, antibiotic-resistant clones were selected by puromycin and then expanded in non-selective medium to be screened by fluorescence microscopy for successful integration of the FUCCI sensor.

### RNA interference

Cells were seeded at a density of 30,000-50,000 cells per well in 6-well plates. The next day, transfection mixes were prepared in sterile DNase-free Eppendorf tubes consisting of 100 µl Opti-MEM (Gibco #11058021), and either 3 µl Mirus TransIT-X2 transfection reagent (Mirus #MIR6000) for hTERT-RPE1 cells, or 3 µl Oligofectamine (Thermo #12252011) for HeLa cells. Solutions were mixed and incubated for 5 min at room temperature, then 60 pM siRNA was added, followed by incubation at room temperature for 30 min, before adding drop-wise to cells. Cells were left to grow for the times indicated in the figure legends. siRNAs used in this study are listed in Table 4.

**Table 4:**
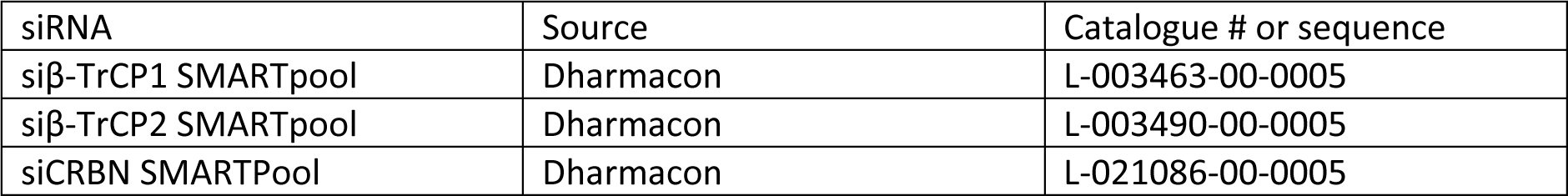
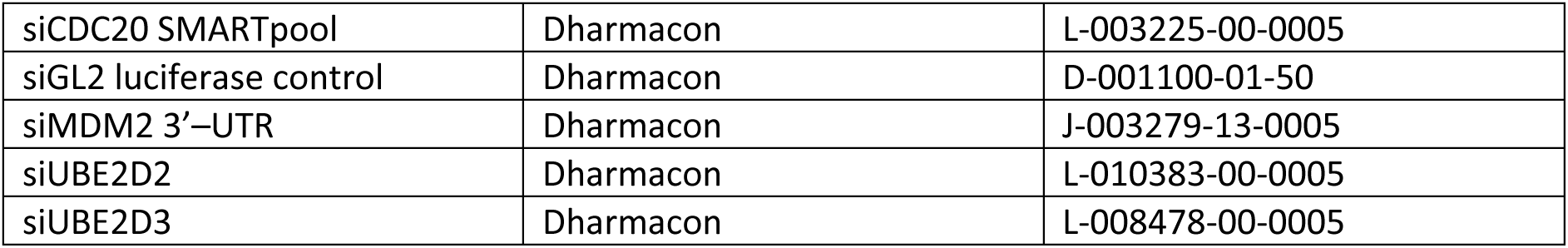
siRNA information for the siRNA sequences used in this study.

| siRNA | Source | Catalogue # or sequence |
| --- | --- | --- |
| siβ-TrCP1 SMARTpool | Dharmacon | L-003463-00-0005 |
| siβ-TrCP2 SMARTpool | Dharmacon | L-003490-00-0005 |
| siCRBN SMARTPool | Dharmacon | L-021086-00-0005 |
| siCDC20 SMARTpool | Dharmacon | L-003225-00-0005 |
| siGL2 luciferase control | Dharmacon | D-001100-01-50 |
| siMDM2 3'-UTR | Dharmacon | J-003279-13-0005 |
| siUBE2D2 | Dharmacon | L-010383-00-0005 |
| siUBE2D3 | Dharmacon | L-008478-00-0005 |

### RNA extraction and cDNA synthesis

Cells were seeded for 70-80% confluency 24 h before lysis. Cells were treated as indicated in the figure legends, washed in PBS, and RNA was extracted using the RNeasy Micro Kit (Qiagen #74004) following the manufacturer’s instructions. RNA purity and yield were assessed on a Nanodrop (Thermo Fisher), and 1-1.5 µg RNA was used a template to make cDNA with the iScript^TM^ cDNA synthesis kit (Bio-rad #1708890), as per the manufacturer’s instructions. cDNA was diluted 1:10 with nuclease-free water before proceeding to RT-qPCR.

### RT-qPCR

Samples were prepared in a 384-well plate format (Bio-rad #HSP3805). Briefly, 4 µl of the diluted cDNA was added to a Master mix containing 4 µl SsoAdvanced Universal SYBR® Green Supermix (Bio-rad #1725270) and 0.5 µl of a 10 µM stock of each Forward and Reverse primer (Table 5) per reaction. Plates were sealed (Thermo Fisher #AB0558) and briefly spun down at 500 *x*g for 30 s. qRT-PCR was performed using a CFX384 Touch Real-Time PCR Detection system (Bio-rad #1855484). All experiments were performed in biological and technical triplicates. Data were quantified using the ΔΔCT method ^33^. PCR amplicons were loaded on to 1% (w/v) agarose gels stained with Midori Green Advance DNA stain (Geneflow #S6-0022) alongside 100 bp ladder (NEB #N3231S), before submitting for DNA Sanger sequencing (Source Genomics, Cambridge, UK) to confirm amplification of the intended targets.

**Table 5:**
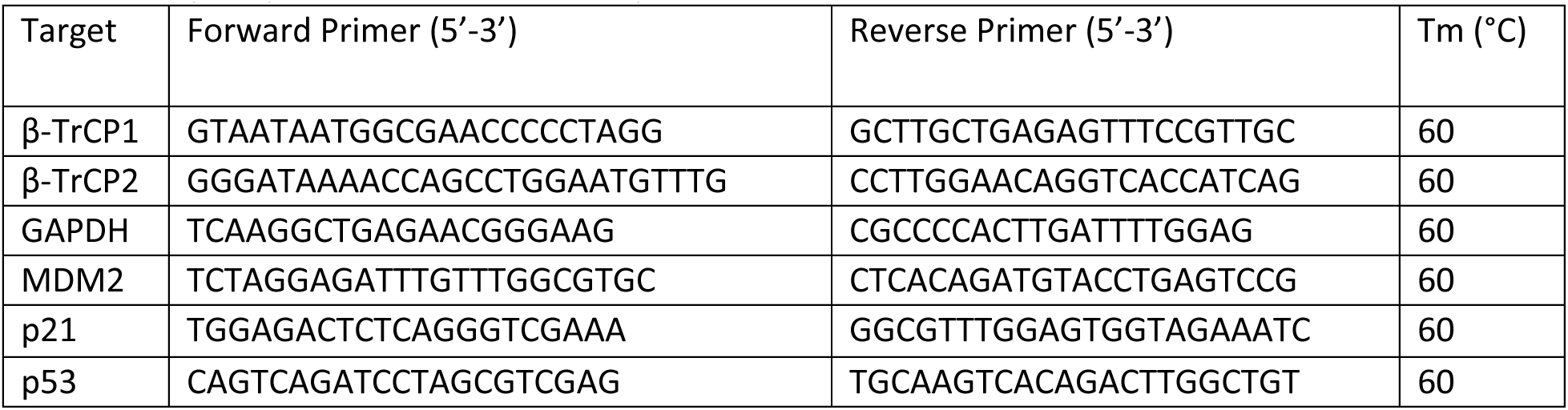
RT-qPCR primers used in this study.

### Mitotic timer assays

For mitotic timer assays, hTERT-RPE1 p53^WT^ or p53^KO^ cells were seeded into 10-cm dishes (five dishes per condition) for 80% confluency the next day. The next day, cells were treated with 25 ng/ml nocodazole for 45 min at 37°C. Mitotic cells were isolated by shake-off, and washed three times in pre-warmed 37°C full medium (with pelleting performed at 200 *x* g for 3 min at 37°C in between washes) before lysing 50% (w/v) of the cells for a mitotic sample, and re-plating the other 50 % (w/v) of the cells for 4 h at 37°C for a G1 sample. Alternatively, following mitotic shake-off, cells were pelleted and re-suspended in full medium containing 25 ng/ml nocodazole and re-plated into 6-cm dishes for longer mitotic samples. Following the longer mitotic arrests, cells were isolated by shake-off, and washed three times in pre-warmed 37°C full medium (with pelleting performed at 200 *x* g for 3 min at 37°C in between washes) before lysing 50% (w/v) of the cells for a mitotic sample, and re-plating the other 50% (w/v) of the cells for 4 h at 37°C for a G1 sample. Where indicated, MD-224 or DMSO (Sigma #D8418) vehicle control was added in the last 45 min of mitosis, before the nocodazole washout procedure. Asynchronous cells were used as controls.

### Cell lysis for Western blots

Following treatments, cells were moved to ice and washed in ice-cold PBS. PBS was removed and cells were lysed in lysis buffer (20 mM Tris-HCl (Sigma-Aldrich, #T4661) pH 7.4, 150 mM NaCl (Sigma-Aldrich #S3014), 1% (v/v) IGEPAL (Sigma-Aldrich #S3014), 0.1% (w/v) sodium deoxycholate (Sigma-Aldrich #D6750), 40 mM sodium β-glycerophosphate (Sigma-Aldrich #G9422), 10 mM NaF (Sigma-Aldrich #201154), 0.3 mM sodium orthovanadate (Sigma-Aldrich #S6508), 100 nM okadaic acid (Enzo Life Sciences #ALX-350-011-M001), 200 nM microcystin-LR (Enzo Life Sciences #ALX-350-012-M001), 1 mM DTT (Sigma-Aldrich #11583786001), 1X protease inhibitor cocktail (Roche, cOmplete^™^, EDTA-free Protease Inhibitor Cocktail, #11873580001), and 1X phosphatase inhibitor cocktail (Sigma, P0044) in water). Lysates were harvested, moved to 1.5-ml Eppendorf tubes (Eppendorf #10451043), and incubated on ice for 10 min. Lysates were then clarified at 14,000 rpm for 15 min at 4°C. Protein concentrations were measured by Bradford assay using Protein Assay Dye Reagent Concentrate (Bio-Rad, 5000006). Samples were normalised to 0.5-1 mg/ml. Sample buffer (0.1875 mM Tris-HCl (Sigma-Aldrich, #T4661) pH 6.8, 1% (w/v) SDS (Sigma-Aldrich #75746), 10% (v/v) glycerol (Sigma-Aldrich #G9012), 0.05% (w/v) bromophenol blue (Sigma-Aldrich #B5525), and 10% (v/v) β-mercaptoethanol (Sigma-Aldrich #M3148) in water) was added, and samples were boiled at 95°C for 10 min, before progressing to SDS-PAGE and Western blotting.

### SDS-PAGE and Western blotting

Proteins were separated by SDS-PAGE (Bio-Rad, Mini-PROTEAN® Tetra cell, #1658000) and transferred onto nitrocellulose membranes (Bio-Rad #1704158, #1704159) using a Trans-blot Turbo system (Bio-Rad #1704150). After transfer, blots were blocked in 5% (w/v) milk powder in PBS (PanReac AppliChem #A0965,9100)-TWEEN® 20 (Sigma-Aldrich #P1379) (PBS-T), before incubation with primary antibodies (Table 1) diluted in 5% (w/v) milk in PBS-T for 1 h at room temperature, or at 4°C overnight. Blots were then washed in PBS-T before incubating with the secondary HRP-conjugated antibodies (Table 1) for 1 h at room temperature, before washing again in PBS-T. Signals were revealed using ECL (GE Healthcare #RPN2106) and visualised on X-ray films (GE Healthcare #GE28-9068-37), developed on an OPTIMAX 2010 X-ray Film Processor (PROTAC-Med).

### Immunoprecipitation

Pellets of 1 × 10^6^ asynchronous or mitotic cells were lysed in 1 ml of lysis buffer (50 mM HEPES pH 7.5, 100 mM NaCl, 0.5% (v/v) Triton X-100, 1 mM DTT, 1:250 Protease Inhibitor Cocktail (Merck #P8340-5ML), 1:1000 Phosphatase Inhibitor cocktail 3 (Merck #P0044-5ML), 100 nM okadaic acid (Enzo LifeSciences #ALX-350-011-M001)) on ice for 30 min, and the lysates were then clarified at 14,000 rpm for 30 min at 4°C. The asynchronous cells were detached by incubating cells in 1 mM EDTA in PBS before pelleting and lysis. The mitotic cells were isolated by shake-off after 100 ng/ml nocodazole treatment for 18 h. 5% (v/v) of the cell lysates were mixed with Sample buffer as input lysates for immunoblotting. The proteins of interest were isolated from 5-20% (v/v) of cell lysates by incubating lysates with 0.5-2.5 µg of antibodies and 20 µl of protein G–Sepharose (Merck, GE17-0618-01) at 4°C for 2 h. The antibody-bound Sepharose beads were then washed three times with lysis buffer at 4°C, resuspended in Sample buffer and boiled at 65°C to elute. IP and input samples were analysed by immunoblotting as described above.

### ^35^S-methionine labelling and immunoprecipitation

Pellets of 2.5 × 10^6^ asynchronous and mitotic cells, isolated as described in the immunoprecipitation section above, were resuspended in 1 ml of pre-warmed labelling medium (Met/Cys/Gln-free DMEM (Thermo, 21013024), containing 0.2 mM L-cysteine, 2 mM GlutaMax (Thermo #35050061), 2.5 mM HEPES (Gibco #15630-056), 1 mM sodium pyruvate (Thermo #11360070), and 10% (v/v) dialysed FBS) with or without 100 ng/ml nocodazole. 200 µCi/ml ^35^S-methionine was added into the cell suspensions for 30 min at 37°C and 5% CO_2_. Following incubation, cells were washed in PBS and pelleted at 200 *x*g for 5 min at 37°C before lysing in 1.25 ml of lysis buffer (50 mM HEPES pH 7.5, 100 mM NaCl, 0.5% (v/v) Triton X-100, 1 mM DTT, 1:250 Protease Inhibitor Cocktail (Merck #P8340-5ML), 1:1000 Phosphatase Inhibitor cocktail 3 (Merck #P0044-5ML), 100 nM okadaic acid (Enzo LifeSciences #ALX-350-011-M001)). Lysates were incubated on ice for 30 min, and then clarified at 14,000 rpm for 30 min at 4°C. An 8% (v/v) aliquot of the cell lysates were mixed with sample buffer to obtain input samples. The proteins of interest were isolated from 8-16% (v/v) of the cell lysates through immunoprecipitation with 1-2.5 µg of antibodies as described in the figures. The proteins were separated by SDS-PAGE, and the gels were fixed and stained using InstantBlue® Coomassie Protein Stain (ISB1L) (Abcam #ab119211). Gels were de-stained with water, and then dried in a gel dryer (BioRad, model #583). Detection of the incorporated label was performed by autoradiography, by exposing gels to x-ray films (GE Healthcare #GE28-9068-37) for 2-7 days at –80°C. Autoradiographs were developed using an OPTIMAX 2010 X-ray Film Processor (PROTAC-Med).

### Cell proliferation assays

To test for proliferation, hTERT-RPE1 p53^WT^ or p53^KO^ cells were treated as described in the mitotic timer assays methods section. Following nocodazole washout, mitotic cells were counted using a Neubauer chamber and were seeded at 5,000 cells per well in 6-well plates. Cells were cultured at 37°C for five days. Following incubation, cells were washed in PBS and fixed/stained in crystal violet/methanol solution (25% (v/v) methanol (VWR #20847.307) + 5% (w/v) crystal violet (Sigma-Aldrich #C0775) in water) for 30 min at room temperature. The crystal violet stain was then removed, and cells washed in water. Plates were left to dry on top of a piece of Whatman paper before imaging on a Bio-Rad Gel Doc XR+ imaging system (Bio-Rad, 1708195EDU). hTERT-RPE1 cells are highly mobile and do not form distinct colonies. The extent of cell proliferation was assessed by densitometry using Fiji/Image J software (National Institutes of Health).

### Cell-cycle tracking using hTERT-RPE1 p53^WT^ and p53^KO^ FUCCI cell lines

For single cell tracking, cells were plated in 35-mm dishes with a 14-mm 1.5 thickness cover glass window on the bottom (MatTek Corp, P35G-1.5-14-C). For imaging, the dishes were placed in a 37°C and 5% CO_2_ environment chamber (Tokai Hit) on the microscope stage. Imaging was performed on an Ultraview Vox spinning disk confocal system running Volocity software (PerkinElmer) using the 20×/0.75 NA UPlanSApo objective on an Olympus IX81 inverted microscope equipped with an electron multiplying charge coupled device (EM-CCD) camera (C9100-13, Hamamatsu Photonics).

Cells were excited by 488 nm and 561 nm lasers with 200 ms exposures at 3-6% laser power. Brightfield reference images were also taken to visualize cell shape with 30 ms exposures. 12-µm image stacks with intervals of 0.6 µm (total 21 planes) were collected at the time intervals indicated in the figures and then maximum intensity projected and cropped in Fiji/Image J (National Institutes of Health) for further analysis. Before the imaging, cells were treated with either 0.1% (v/v) DMSO as a control, 25 ng/ml (82.5 nM) nocodazole, 30 nM CENP-E inhibitor, 100 nM MD-224, 2.5 µM Nutlin-3A or combinations of these chemicals for 30 min or 4 hours at 37°C, 5% CO_2_ and then washed five times in pre-warmed 37°C full medium. After the washout, the dish was immediately placed on the microscope, and then the mitotic cells expressing mAG-hGem, in prometaphase or metaphase, were imaged and tracked to measure G1 duration.

### Immunofluorescence microscopy

Cells plated on glass coverslips were washed in PBS, and fixed with 3% (w/v) PFA (Sigma-Aldrich #158127) in PBS pH 7.4 at room temperature for 15 min. Unreacted PFA was quenched by washing with 50 mM NH_4_Cl (Sigma-Aldrich #213330) in PBS, following by a 10-min incubation in 50 mM NH_4_Cl in PBS. Cells were washed three times in PBS, followed by permeabilization with 0.2% (v/v) Triton^TM^ X-100 (Sigma-Aldrich #X100) in PBS at room temperature for 5-10 min. Permeabilised cells were washed three times in PBS before incubating with the appropriate primary antibodies (Table 1). Antibody dilutions were performed in PBS, and coverslips were placed cell-side down on droplets of antibody solutions, inside a humidified chamber for 1 h at room temperature. Following incubation, cells were washed three times in PBS, and incubated with the secondary AlexaFluor-conjugated antibodies (Table 1) for 1 h at room temperature, as described for the primary antibody incubation. DAPI (Sigma-Aldrich #D9542) was added at the same time as the secondary antibodies, in order to stain DNA. Following incubation, cells were washed three times in PBS, and once in water, before mounting onto glass microscopy slides with Mowiol® 4-88 (Sigma-Aldrich #81381). Samples were imaged on a standard upright microscope system (BX61, Olympus) with filter sets for DAPI, GFP/Alexa Fluor 488, Cy3/Alexa Fluor 555 and Cy5/Alexa Fluor 647 (Chroma Technology Corp.), a 2048 × 2048-pixel complementary metal oxide semiconductor camera (Prime, Photometrics), and MetaMorph 7.5 imaging software (Molecular Devices). Illumination was provided by an LED light source (pE300, CoolLED Illumination Systems). Image stacks with a spacing of 0.4 µm through the cell volume were maximum intensity projected and cropped in Image J (National Institutes of Health).

### Statistical analysis

Statistical analysis was performed using GraphPad Prism 9.3.1 (GraphPad Software). Statistical significance was analysed using an unpaired two-tailed t-test with Welch’s correction, a one-way ANOVA, a Brown-Forsythe ANOVA, a Kruskal-Wallis test, or Dunn’s multiple comparison test. Graphs display the mean ± SD or SEM. P-values are shown on graphs as follows: p ≥ 0.05=not significant (ns), p < 0.05=*, p < 0.01=**, p < 0.001=***, p < 0.0001=****.

## Notes

### Competing Interest Statement

The authors have declared no competing interest.

### Summary of Updates

In data shown in the new Figure 1, we demonstrate that the mitotic timer pathway responds to the normal stochastic variation in the length of mitosis with a p53-dependent arrest in G1. We show that this response, and its increased frequency under specific stress conditions, is explained by the specific biochemical situation found in mitosis: attenuation of transcription and translation combined with self-catalysed destruction of the p53 ubiquitin-ligase MDM2, which enables the cell to use MDM2 levels as a precise read-out of mitotic duration. To strengthen this conclusion, we have measured MDM2 synthesis in mitosis, and directly confirmed a key prediction of our hypothesis that this is reduced <95% in mitotic cells (new Figure 2). Using a combination of biochemical analysis and single cell-imaging of both FUCCI and p21-GFP cell lines we determine the critical threshold for MDM2 in mitosis, below which it becomes limiting for p53 regulation at the onset of G1 leading to rapid p21 induction and cell cycle arrest (Figure 4-5 and new Figure 6).

## References

1 Wong, Y. L. et al. Cell biology. Reversible centriole depletion with an inhibitor of Polo-like kinase 4. Science 348, 1155–1160, doi:10.1126/science.aaa5111 (2015).

2 Thompson, S. L. & Compton, D. A. Proliferation of aneuploid human cells is limited by a p53-dependent mechanism. J Cell Biol 188, 369–381, doi:10.1083/jcb.200905057 (2010).

3 Li, R. & Zhu, J. Effects of aneuploidy on cell behaviour and function. Nat Rev Mol Cell Biol 23, 250–265, doi:10.1038/s41580-021-00436-9 (2022).

4 Thompson, S. L. & Compton, D. A. Examining the link between chromosomal instability and aneuploidy in human cells. J Cell Biol 180, 665–672, doi:10.1083/jcb.200712029 (2008).

5 Uetake, Y. & Sluder, G. Prolonged prometaphase blocks daughter cell proliferation despite normal completion of mitosis. Curr Biol 20, 1666–1671, doi:10.1016/j.cub.2010.08.018 (2010).

6 Yang, Z., Loncarek, J., Khodjakov, A. & Rieder, C. L. Extra centrosomes and/or chromosomes prolong mitosis in human cells. Nat Cell Biol 10, 748–751, doi:10.1038/ncb1738 (2008).

7 Lambrus, B. G. et al. A USP28-53BP1-p53-p21 signaling axis arrests growth after centrosome loss or prolonged mitosis. J Cell Biol 214, 143–153, doi:10.1083/jcb.201604054 (2016).

8 Meitinger, F. et al. 53BP1 and USP28 mediate p53 activation and G1 arrest after centrosome loss or extended mitotic duration. J Cell Biol 214, 155–166, doi:10.1083/jcb.201604081 (2016).

9 Cuella-Martin, R. et al. 53BP1 Integrates DNA Repair and p53-Dependent Cell Fate Decisions via Distinct Mechanisms. Mol Cell 64, 51–64, doi:10.1016/j.molcel.2016.08.002 (2016).

10 Fong, C. S. et al. 53BP1 and USP28 mediate p53-dependent cell cycle arrest in response to centrosome loss and prolonged mitosis. Elife 5, doi:10.7554/eLife.16270 (2016).

11 Sakaue-Sawano, A. et al. Visualizing spatiotemporal dynamics of multicellular cell-cycle progression. Cell 132, 487–498, doi:10.1016/j.cell.2007.12.033 (2008).

12 Kubbutat, M. H., Jones, S. N. & Vousden, K. H. Regulation of p53 stability by Mdm2. Nature 387, 299–303, doi:10.1038/387299a0 (1997).

13 Haupt, Y., Maya, R., Kazaz, A. & Oren, M. Mdm2 promotes the rapid degradation of p53. Nature 387, 296–299, doi:10.1038/387296a0 (1997).

14 Honda, R., Tanaka, H. & Yasuda, H. Oncoprotein MDM2 is a ubiquitin ligase E3 for tumor suppressor p53. FEBS Lett 420, 25–27, doi:10.1016/s0014-5793(97)01480-4 (1997).

15 Blackford, A. N. & Jackson, S. P. ATM, ATR, and DNA-PK: The Trinity at the Heart of the DNA Damage Response. Mol Cell 66, 801–817, doi:10.1016/j.molcel.2017.05.015 (2017).

16 Fang, S., Jensen, J. P., Ludwig, R. L., Vousden, K. H. & Weissman, A. M. Mdm2 is a RING finger-dependent ubiquitin protein ligase for itself and p53. J Biol Chem 275, 8945–8951, doi:10.1074/jbc.275.12.8945 (2000).

17 Inuzuka, H. et al. Phosphorylation by casein kinase I promotes the turnover of the Mdm2 oncoprotein via the SCF(beta-TRCP) ubiquitin ligase. Cancer Cell 18, 147–159, doi:10.1016/j.ccr.2010.06.015 (2010).

18 Fan, H. & Penman, S. Regulation of protein synthesis in mammalian cells. II. Inhibition of protein synthesis at the level of initiation during mitosis. J Mol Biol 50, 655–670, doi:10.1016/0022-2836(70)90091-4 (1970).

19 Taylor, J. H. Nucleic acid synthesis in relation to the cell division cycle. Ann N Y Acad Sci 90, 409–421, doi:10.1111/j.1749-6632.1960.tb23259.x (1960).

20 Prescott, D. M. & Bender, M. A. Synthesis of RNA and protein during mitosis in mammalian tissue culture cells. Exp Cell Res 26, 260–268, doi:10.1016/0014-4827(62)90176-3 (1962).

21 Blagosklonny, M. V. Prolonged mitosis versus tetraploid checkpoint: how p53 measures the duration of mitosis. Cell Cycle 5, 971–975, doi:10.4161/cc.5.9.2711 (2006).

22 Santaguida, S. & Amon, A. Short-and long-term effects of chromosome mis-segregation and aneuploidy. Nat Rev Mol Cell Biol 16, 473–485, doi:10.1038/nrm4025 (2015).

23 Scheffner, M., Huibregtse, J. M., Vierstra, R. D. & Howley, P. M. The HPV-16 E6 and E6-AP complex functions as a ubiquitin-protein ligase in the ubiquitination of p53. Cell 75, 495–505, doi:10.1016/0092-8674(93)90384-3 (1993).

24 Barak, Y., Juven, T., Haffner, R. & Oren, M. mdm2 expression is induced by wild type p53 activity. EMBO J 12, 461–468, doi:10.1002/j.1460-2075.1993.tb05678.x (1993).

25 Tanenbaum, M. E., Stern-Ginossar, N., Weissman, J. S. & Vale, R. D. Regulation of mRNA translation during mitosis. Elife 4, doi:10.7554/eLife.07957 (2015).

26 Krenning, L., Sonneveld, S. & Tanenbaum, M. E. Time-resolved single-cell sequencing identifies multiple waves of mRNA decay during the mitosis-to-G1 phase transition. Elife 11, doi:10.7554/eLife.71356 (2022).

27 Nomura, K. et al. Structural analysis of MDM2 RING separates degradation from regulation of p53 transcription activity. Nat Struct Mol Biol 24, 578–587, doi:10.1038/nsmb.3414 (2017).

28 Saville, M. K. et al. Regulation of p53 by the ubiquitin-conjugating enzymes UbcH5B/C in vivo. J Biol Chem 279, 42169–42181, doi:10.1074/jbc.M403362200 (2004).

29 Nakayama, K. I. & Nakayama, K. Ubiquitin ligases: cell-cycle control and cancer. Nat Rev Cancer 6, 369–381, doi:10.1038/nrc1881 (2006).

30 Li, Y. et al. Discovery of MD-224 as a First-in-Class, Highly Potent, and Efficacious Proteolysis Targeting Chimera Murine Double Minute 2 Degrader Capable of Achieving Complete and Durable Tumor Regression. J Med Chem 62, 448–466, doi:10.1021/acs.jmedchem.8b00909 (2019).

31 Burigotto, M. et al. PLK1 promotes the mitotic surveillance pathway by controlling cytosolic 53BP1 availability. EMBO Rep, e57234, doi:10.15252/embr.202357234 (2023).

32 Barr, A. R. et al. DNA damage during S-phase mediates the proliferation-quiescence decision in the subsequent G1 via p21 expression. Nat Commun 8, 14728, doi:10.1038/ncomms14728 (2017).

33 Livak, K. J. & Schmittgen, T. D. Analysis of relative gene expression data using real-time quantitative PCR and the 2(-Delta Delta C(T)) Method. Methods 25, 402–408, doi:10.1006/meth.2001.1262 (2001).

